# CRISPR/Cas9 screen identifies KRAS-induced COX-2 as a driver of immunotherapy resistance in lung cancer

**DOI:** 10.1101/2023.04.13.536740

**Authors:** Jesse Boumelha, Andrea de Castro, Nourdine Bah, Hongui Cha, Sophie de Carné Trécesson, Sareena Rana, Panayiotis Anastasiou, Edurne Mugarza, Christopher Moore, Robert Goldstone, Phil East, Kevin Litchfield, Se-Hoon Lee, Miriam Molina-Arcas, Julian Downward

## Abstract

Oncogenic KRAS impairs anti-tumour immune responses, but effective strategies to combine KRAS inhibitors and immunotherapies have so far proven elusive. In vivo CRISPR-Cas9 screening in an immunogenic murine lung cancer model identifies mechanisms by which oncogenic KRAS promotes immune evasion, most notably expression of immunosuppressive cyclooxygenase-2 (COX-2) in cancer cells. Oncogenic KRAS was a potent inducer of COX-2 in both mouse and human lung cancer which was suppressed using KRAS inhibitors. COX-2 acting via prostaglandin E2 (PGE2) promotes resistance to immune checkpoint blockade (ICB) in both mouse and human lung adenocarcinoma. Targeting COX-2/PGE2 remodelled the tumour microenvironment by inducing pro-inflammatory polarisation of myeloid cells and influx of activated cytotoxic CD8+ T cells, which increased the efficacy of ICB. Restoration of COX-2 expression contributed to tumour relapse after prolonged KRAS inhibition. We propose testing COX-2/PGE2 pathway inhibitors in combination with KRAS G12C inhibition or ICB in patients with KRAS-mutant lung cancer.

## INTRODUCTION

Despite improvements in systemic therapies, lung adenocarcinoma (LUAD) remains the most common cause of cancer-related deaths worldwide (1). Immune checkpoint blockade (ICB), which can reinvigorate anti-tumour immunity, has shown remarkable clinical success in multiple cancer types (2), including LUAD (3), achieving durable responses in a subset of patients. Monoclonal antibodies targeting the immunosuppressive PD-L1/PD-1 axis have become standard of care for LUAD patients, either as a monotherapy (4) or in combination with chemotherapy (5). However, only a fraction of patients benefits from ICB highlighting the need for combination strategies that will broaden responses to current immunotherapies. Recent clinical efforts, such as the SKYSCRAPER-01 trial combining PD-L1 blockade with an anti-TIGIT antibody, have failed to lead to improved responses in lung cancer and there is a need to better understand mechanisms of immune evasion in order to design rational combination strategies that are more likely to provide clinical benefit.

Mutations in the oncogene KRAS drive tumourigenesis in 30% of LUAD cases (6). However, the development of inhibitors that directly target KRAS has been notoriously challenging (7). A major breakthrough was achieved with the recent development of mutant-specific KRAS^G12C^ inhibitors (8) which covalently bind to the novel cysteine residue present in nearly half of all KRAS-mutant LUAD patients (9). This has led to the approval of Amgen’s clinical compound sotorasib for the treatment of locally advanced or metastatic KRAS^G12C^-mutant lung cancer (10). Whilst these drugs have mild toxicities and achieve clinical responses in a substantial proportion of patients, as with other targeted therapies, responses are often short-lived as resistance inevitably arises (11).

Accumulating evidence suggests that oncogenic signalling extends beyond the tumour cell compartment and engages with the host stromal and immune compartments. Consequently, oncogenic drivers have been shown to play dominant roles in shaping the tumour immune landscape of different cancers and inhibiting anti-tumour immune responses (12). Analysis of clinical samples has demonstrated that KRAS mutations are associated with an immunosuppressive tumour microenvironment (TME) (13) and preclinical studies have identified a number of mechanisms by which oncogenic KRAS can drive immune evasion including promoting the expression of numerous immunosuppressive myeloid chemoattractants (14,15) and immune checkpoint ligands (16). Together these observations provide a rational basis for combining KRAS inhibitors with ICB and numerous preclinical studies have demonstrated that this combination leads to improved therapeutic responses, at least in immunogenic models (15,17,18). However, recent reports of the CODEBREAK 100/101 clinical trial evaluation of sotorasib in combination with anti-PD-L1/PD-1 antibodies have shown serious toxicities (19,20), casting doubt on the viability of this combination. As an alternative approach, a greater understanding of how oncogenic KRAS drives immune evasion may identify novel immunotherapy combination strategies that could improve outcomes for KRAS-mutant lung cancer patients.

Pooled CRISPR screens have been increasingly used to uncover tumour-intrinsic determinants of anti-tumour immunity, identifying numerous genes that either promote sensitivity or resistance to immune control (21–25). We recently developed a novel immunogenic model of KRAS-mutant lung cancer allowing for the preclinical study of tumour-immune interactions (17). Here we carry out a pooled *in vivo* CRISPR screen in this novel model to interrogate the role of 240 KRAS-regulated genes in controlling anti-tumour immunity. This identified several genes that increased sensitivity or resistance to anti-tumour immune responses. Among these, the prostaglandin synthase cyclooxygenase-2 (COX-2), responsible for synthesis of the immunosuppressive molecule prostaglandin E2 (PGE_2_), was identified as a major driver of immune evasion and resistance to immunotherapy. Targeting of the COX2/PGE_2_ axis improved the response of KRAS-mutant lung tumours to anti-PD-1 therapy by inducing pro-inflammatory polarisation of myeloid cells and enhancing T cell infiltration and activation. Importantly, oncogenic KRAS signalling was a strong driver of the COX-2/PGE_2_ signalling axis in both mouse and human LUAD and COX-2 inhibition delayed tumour relapse after KRAS^G12C^ inhibition.

## RESULTS

### *In vivo* CRISPR screen identifies tumour-intrinsic determinants of anti-tumour immunity

It is challenging to carry out large scale genome-wide screens *in vivo*, whilst also maintaining a sufficiently high representation of the pooled library, as there is a limit on the number of cells that can be orthotopically transplanted. Instead, a rationally selected, smaller, customised library was used. First, we generated a library of lentiviral vectors encoding sgRNAs targeting 240 genes that are regulated by KRAS in human LUAD (Supplementary Table S1). KRAS-regulated genes were identified by differential gene expression analysis of TCGA LUAD samples and LUAD cell lines from the Cancer Cell Line Encyclopedia (CCLE) which were stratified as having high or low RAS pathway activity using a novel RAS transcriptional signature (26). Additional genes were identified using RNA-seq data from KRAS^G12C^-mutant LUAD cell lines (H358 and H23) treated with a KRAS inhibitor and immortalised type II pneumocytes expressing an ER-KRAS^G12V^ fusion protein which can be readily activated by administration of 4-hydroxytamoxifen (4-OHT) (15). To carry out the screen we utilised the immunogenic KPAR cell line, derived from a genetic KRAS^G12D^ p53^−/−^ lung cancer mouse model as it stimulates endogenous anti-tumour immune responses and is partially responsive to immunotherapy (17). Next, we engineered the immunogenic KRAS^G12D^ p53^−/−^ KPAR cell line to express Cas9 under a doxycycline-inducible promoter (Supplementary Fig. 1A). An inducible system was chosen as it allows temporal control of editing and circumvents any compounding consequences of Cas9 immunogenicity *in vivo*. Importantly, Cas9 expression was abrogated *in vitro* 48 hours after the removal of doxycycline (Supplementary Fig. 1B) and was not re-expressed *in vivo* (Supplementary Fig. 1C).

Library-transduced KPAR iCas9 cells were treated with doxycycline for four days to allow gene-editing to occur followed by a two-day washout period to abrogate the expression of Cas9 before orthotopic transplantation into C57BL/6 (WT) mice or *Rag2^−/−^*;*Il2rg^−/−^* mice which lack T cells, B cells and NK cells and therefore do not exert anti-tumour immune responses (Fig. 1A). After three weeks, genomic DNA was isolated from tumour-bearing lungs and subject to deep next-generation sequencing (NGS) to compare library representation in tumours growing in immune-competent and immune-deficient mice. In parallel, NGS of genomic DNA from cells passaged *in vitro* was carried out to identify genes that affect cell viability. Analysis of genes targeted by sgRNAs that were depleted *in vitro* identified a number of genes known to affect cell viability in KRAS-mutant tumour cells, including *Myc* (27) and *Fosl1* (28), thereby validating the functionality of the KRAS-target sgRNA library (Fig. 1B). Furthermore, a number of sgRNAs were equally depleted in immune-competent and immune-deficient mice compared to *in vitro* passaged cells, including those targeting the EMT regulator *Zeb1* and the anti-apoptotic caspase inhibitor c-FLIP (encoded by *Cflar*) (Fig. 1C). These genes therefore supported tumour growth *in vivo* by mechanisms independent of anti-tumour immunity.

**Figure 1.**
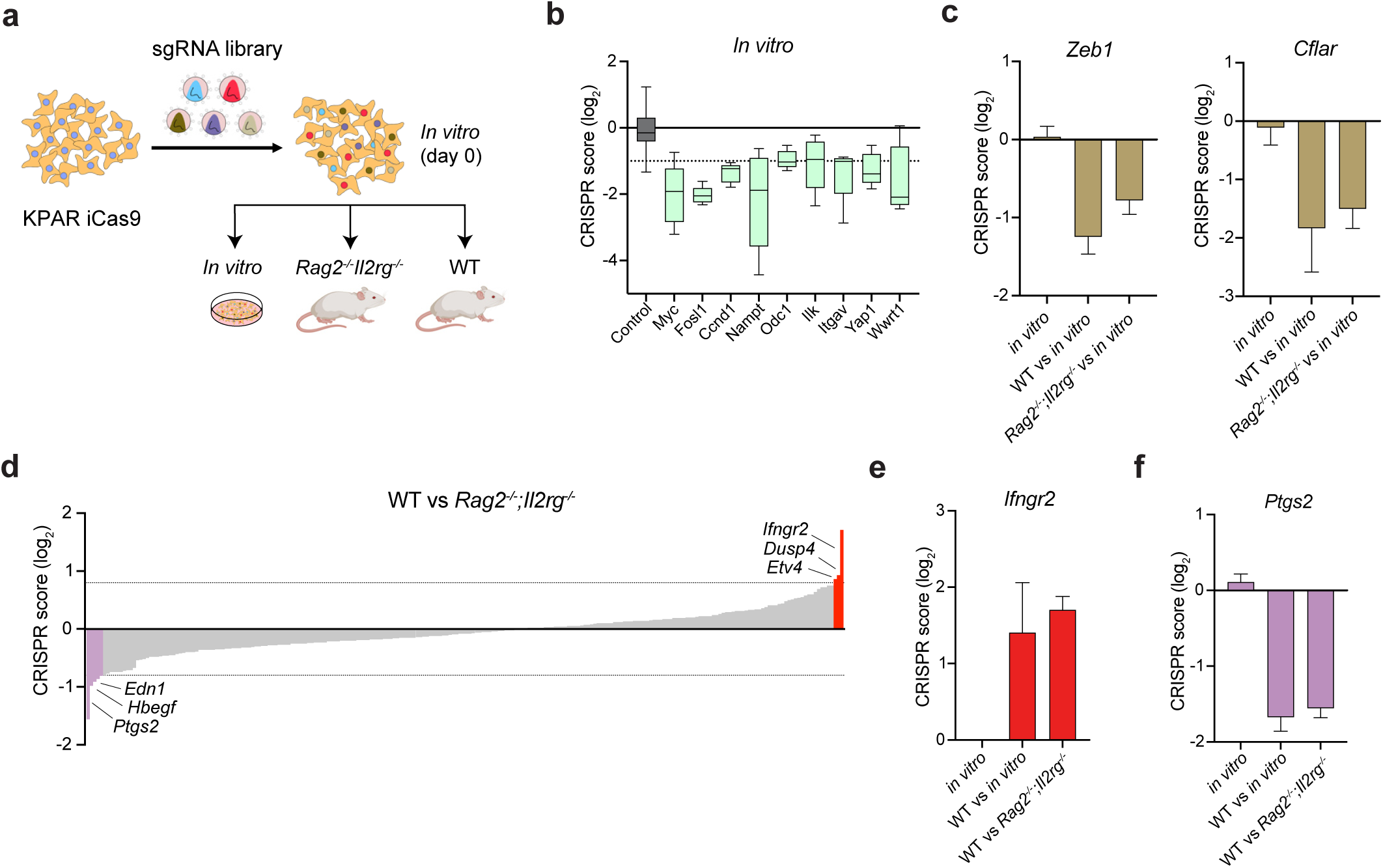
In vivo CRISPR-Cas9 screen identifies regulators of anti-tumour immunity. (A) Schematic of pooled CRISPR-Cas9 screen. (B) sgRNAs targeting genes depleted in vitro compared to non-target controls. CRISPR score is defined as the average log2-fold change in abundance of sgRNA reads at day 28 (in vitro) vs day 0 (in vitro) for each gene. (C) sgRNAs targeting Cflar and Zeb1 depleted in vivo in immune-competent and immune-deficient mice. (D) Average log2-fold change in abundance of sgRNA reads for all genes in immune-competent (WT) versus Rag2-/-;Il2rg-/- mice. (E-F) Enrichment of sgRNAs targeting Ifngr2 (E) and depletion of sgRNAs targeting Ptgs2 (F) in WT versus Rag2-/-;Il2rg-/- mice. Data are mean ± SEM for (C and E-F).

By comparing sgRNAs that were depleted or enriched in immune-competent compared to immune-deficient mice we identified genes that modulate anti-tumour immune responses (Fig. 1D). The most enriched sgRNAs in immune-competent mice were targeting a subunit of the IFNγ receptor (*Ifngr2)* (Fig. 1E). Furthermore, we also saw enrichment of sgRNAs targeting the Ets transcription factor ETV4 (Supplementary Fig. 1D). Importantly, *Etv4^−/−^* KPAR tumours grew faster than parental tumours in WT mice but not in *Rag2^−/−^*;*Il2rg^−/−^*mice (Supplementary Fig. 1E), confirming a role for ETV4 in sensitising tumours to anti-tumour immune responses. Indeed, gene expression analysis of *Etv4^−/−^* KPAR tumours demonstrated that loss of ETV4 resulted in a drastic downregulation of multiple anti-tumour immunity genes (Supplementary Fig. 1F). Conversely, the most depleted sgRNAs in immune-competent mice targeted the prostaglandin synthase COX-2, encoded by *Ptgs2* (Fig. 1F). In addition, sgRNAs targeting the secreted protein endothelin-1, encoded by *Edn1*, were also depleted in immune-competent mice (Supplementary Fig. 1G). Confirming the role of tumour-derived EDN1 in facilitating immune evasion, *Edn1^−/−^* KPAR tumours grew slower than parental tumours specifically in WT mice (Supplementary Fig. 1H).

### KRAS suppresses anti-tumour immunity by inhibition of tumour-intrinsic IFNγ signalling

Tumour-intrinsic IFNγ signalling has previously been shown to be required for anti-tumour immune responses in carcinogen-induced mouse models of cancer (29) and responses to immunotherapy in melanoma (30). To validate the role of tumour-intrinsic IFNγ signalling in anti-tumour immune responses uncovered in the screen (Fig. 1E) we generated *Ifngr2*^−/−^ KPAR cell lines using CRISPR-Cas9. Flow cytometry confirmed that *Ifngr2*^−/−^ KPAR cells had lost the expression of the IFNγ-receptor β subunit (Supplementary Fig. 2A). As expected, *Ifngr2*^−/−^ KPAR cells were insensitive to IFNγ *in vitro* as we were unable to detect pSTAT1 in response to IFNγ (Supplementary Fig. 2B). Whilst *Ifngr2*^−/−^ cells grew similarly to parental cells *in vitro* (Supplementary Fig. 2C), they grew faster than parental cells when transplanted into immune-competent mice (Fig. 2A). In contrast *Ifngr2*^−/−^ and parental tumours grew at similar rates in immune-deficient *Rag2^−/−^*;*Il2rg^−/−^*mice. These data were validated using a second clone (Supplementary Fig. 2D) indicating that intact tumour-intrinsic IFNγ signalling is required for effective anti-tumour immunity.

**Figure 2.**
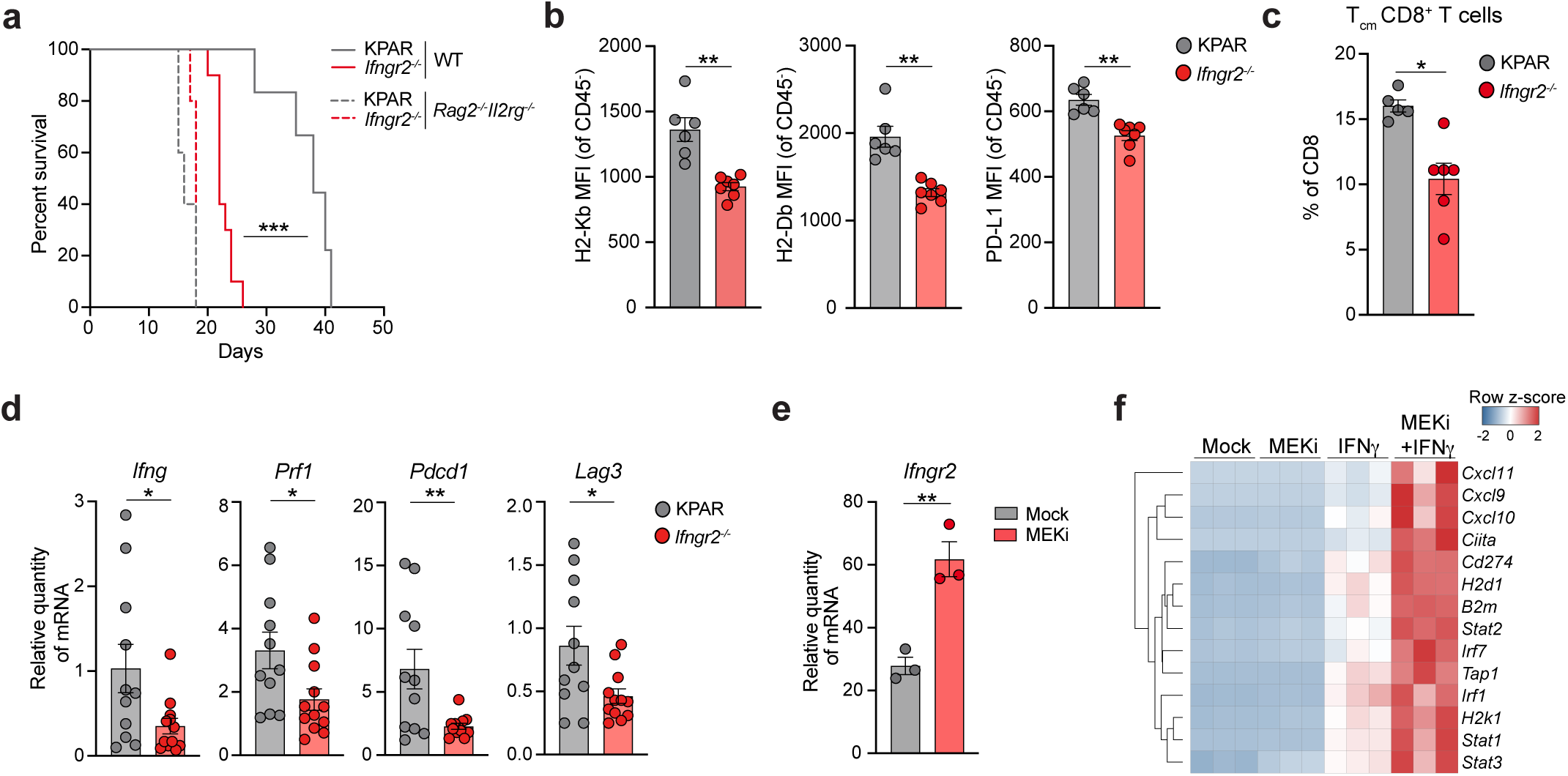
KRAS-driven inhibition of tumour-intrinsic IFN signalling promotes immune evasion. (A) Kaplan-Meier survival of immune-competent or *Rag2^−/−^;Il2rg^−/−^* mice following orthotopic transplantation with KPAR cells or *Ifngr2^−/−^* cells, n=5-10 per group. Analysis of survival curves was carried out using log-rank (Mantel-Cox) test; *** P<0.001. (B) Flow cytometry analysis showing surface expression of H2-Kb, H2-Db and PD-L1 on CD45-cells in KPAR and *Ifngr2^−/−^* tumours. (C) mRNA expression by qPCR of immune-related genes. (D) mRNA expression of Ifngr2 in KPAR cells treated with 10nM trametinib for 24h. (E) Heatmap showing expression of IFN-response genes by qPCR in KPAR cells treated for 24h with 100ng/ml recombinant IFNγ, 10nM trametinib or a combination of both. Data are mean ± SEM for (B-D), n=6-11 per group (B-C). Groups were compared using unpaired, two-tailed Student’s t-test; * P<0.05, ** P<0.01.

IFNγ has been shown to have direct anti-proliferative effects in human cancer cell lines (30). However, the growth of KPAR cells *in vitro* was unaffected by treatment with IFNα, IFNβ or IFNγ (Supplementary Fig. 2E). Tumour-intrinsic IFNγ signalling also regulates the expression of antigen presentation machinery and T cell chemoattractants. *Ifngr2*^−/−^ KPAR cells failed to upregulate IFN signalling molecules (*Irf1, Stat1*), IFN-response genes (*Cxcl9, Cxcl10, Cd274*) and antigen presentation machinery genes (*B2m, Tap1, H2d1, H2k1*) upon treatment with IFNγ (Supplementary Fig. 2F). Furthermore, *Ifngr2*^−/−^ KPAR cells were unable to upregulate PD-L1 or MHC-I surface expression in response to IFNγ, but remained sensitive to type I (IFNα and IFNβ) interferons (Supplementary Fig. 2G). Consistent with this, CD45^−^ cells, which includes tumour and stromal cells, had reduced surface expression of MHC-I and PD-L1 in *Ifngr2*^−/−^ KPAR tumours (Fig. 2B, Supplementary Fig. 2H). In agreement with the hypothesis that tumour-intrinsic IFNγ signalling is required anti-tumour immunity, *Ifngr2*^−/−^ KPAR tumours had less central memory (CD62L^+^CD44+) CD8^+^ T cells which are important for durable anti-tumour immune responses (Fig. 2C). Furthermore, several genes involved in cytotoxic anti-tumour immune responses were downregulated in *Ifngr2*^−/−^ tumours including those encoding IFNγ, perforin, and the checkpoint molecules PD-1 and LAG-3 (Fig. 2D).

Previous studies have demonstrated that oncogenic KRAS regulates tumour-intrinsic IFN pathway gene expression (15). Indeed, treatment of KPAR cells with the MEK inhibitor trametinib (MEKi), or a previously published three-drug KRAS pathway inhibitor combination (31) of trametinib, everolimus and linsitinib (TEL), resulted in upregulation of the IFNγ-receptor β subunit (Fig. 2D, Supplementary Fig. 2I). Moreover, the transcriptional upregulation of IFN-response genes in KPAR cells treated *in vitro* with IFNγ was greatly potentiated by MEKi treatment, validating previous results in other lung cancer models (Fig. 2E). Oncogenic KRAS similarly supressed the response of KPAR cells to IFNα (Supplementary Fig. 2J). The oncogene Myc acts as a transcriptional suppressor of type I IFN-response genes in pancreatic cancer (32) and we have recently shown that Myc mediates KRAS-driven inhibition of IFN-response genes in lung cancer (15). Myc expression is often driven by KRAS and treatment of KPAR cells with MEKi led to rapid downregulation of Myc (Supplementary Fig. 2K). Consistent with its role in suppressing IFN-responses we saw increased expression of IFN-genes in Myc-depleted cells treated with IFNγ (Supplementary Fig. 2L). Furthermore, Myc-depletion led to increased expression of IFN signalling molecules (STAT1 and STAT2) which were not further upregulated upon treatment with MEKi (Supplementary Fig. 2M), suggesting that oncogenic KRAS suppresses tumour-intrinsic IFN responses by driving expression of Myc. In summary, these data suggest that KRAS-mediated inhibition of tumour-intrinsic IFNγ responses, which is required for effective anti-tumour immunity, may contribute to immune evasion in KRAS-mutant lung cancer.

### Tumour-intrinsic COX-2 suppresses innate and adaptive anti-tumour immunity

*Ptgs2* loss was the strongest sensitiser to anti-tumour immunity in the screen and encodes the enzyme COX-2 which is overexpressed in many cancer types. COX-2 is responsible for the synthesis of the prostanoid PGE_2_, which has been shown to suppress anti-tumour immunity in preclinical models of colorectal cancer and melanoma (33). To validate the results obtained in the screen (Fig. 1F), we began by generating *Ptgs2*^−/−^ KPAR cell lines using CRISPR-Cas9. Western blotting confirmed the loss of COX-2 expression and PGE_2_ production in *Ptgs2*^−/−^ cells (Supplementary Fig. 3A). We did not observe any difference in the growth of *Ptgs2*^−/−^ cells and parental cells *in vitro* (Supplementary Fig. 3B). However, they grew considerably slower when orthotopically transplanted into immune-competent mice, which as a result had significantly increased survival, with 60% of mice experiencing complete rejection (Fig. 3A). Importantly, COX-2-deficient tumours grew similarly to parental tumours when transplanted into immune-deficient *Rag2^−/−^*;*Il2rg^−/−^*mice. This was further validated using a second clone (Supplementary Fig. 3C). Therefore, *Ptgs2*^−/−^ cell lines showed no cell-autonomous defects in tumour progression but were instead sensitised to anti-tumour immune responses, with immunological rejection occurring in a substantial proportion of mice.

**Figure 3.**
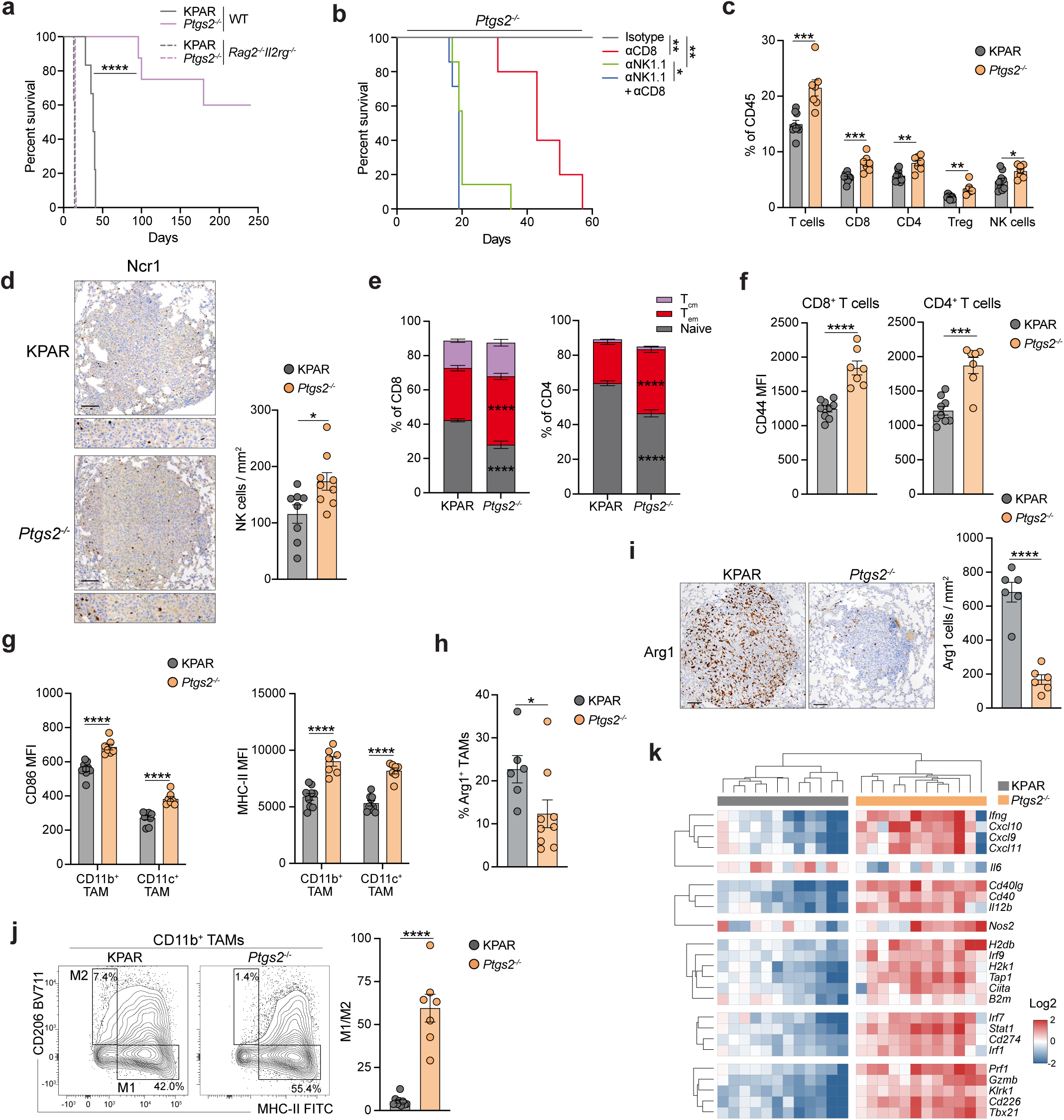
Tumour-intrinsic COX-2 suppresses anti-tumour immunity. (A) Kaplan-Meier survival of immune-competent or *Rag2^−/−^;Il2rg^−/−^* mice following orthotopic transplantation with KPAR cells or *Ptgs2^−/−^* cells (n=5-10 per group). Analysis of survival curves was carried out using log-rank (Mantel-Cox) test; **** P<0.0001. (B) Kaplan-Meier survival of mice treated with 200µg anti-NK1.1 and/or 200µg anti-CD8 or corresponding isotype control (n=5-7 per group) after orthotopic transplantation of *Ptgs2^−/−^* cells. Treatment was initiated 1 day before transplantation and was administered once weekly until endpoint. Analysis of survival curves was carried out using log-rank (Mantel-Cox) test; * P<0.05, ** P<0.01. (C) Frequency of tumour-infiltrating T cell populations and NK cells in KPAR and *Ptgs2^−/−^* orthotopic tumours. (D) Quantification and representative immunohistochemistry staining for NKp46^+^ NK cells. Scale bar represents 100µm. (E) Stacked bar plots showing frequency of central memory (CD62L^+^CD44^+^), effector memory (CD44^+^CD62L^−^) and naïve (CD62L^−^CD44^−^) CD8^+^ (left) and CD4^+^ (right) T cells. (F) Surface expression of CD44 on CD8^+^ (left) and CD4^+^ (right) T cells. (G) Surface expression of CD86 (left) and MHC-II (right) on CD11b^+^ macrophages and CD11c^+^ macrophages. (H) Percentage of Arg1^+^ CD11b^+^ macrophages. (I) Quantification and representative immunohistochemistry staining for the immunosuppressive macrophage marker Arg1. Scale bar represents 100µm. (J) Representative flow cytometry plots of CD206 and MHC-II surface expression on CD11b^+^ macrophages (left) and quantification of M1/M2 ratio based on the gated populations (right). (K) Heatmap showing hierarchical clustering of KPAR and Ptgs2-/- tumours based on mRNA expression of anti-tumour immunity genes assessed by qPCR. Data are mean ± SEM for (C-J), n=6-9 per group. Symbols represent pooled tumours from individual mice. Statistics were calculated by paired, two-tailed Student’s t-test (C-D and F-J) or two-way ANOVA, FDR 0.05 (E); * P<0.05, ** P<0.01, *** P<0.001, **** P<0.0001.

As *Rag2^−/−^*;*Il2rg^−/−^* mice lack NK cells, T cells and B cells, we wanted to decipher the contribution of innate and adaptive immunity to the reduced growth of COX-2-deficient tumours in immune-competent mice. COX-2-deficient tumours grew faster in mice treated with antibodies depleting either NK cells or CD8^+^ T cells and grew fastest in mice lacking both subsets (Fig. 3B). Interestingly, tumours grew faster in mice lacking NK cells compared to mice lacking CD8^+^ T cells, suggesting the innate immune response was largely responsible for the impaired growth of COX-2-deficient tumours, as previously reported (34). However, no mice survived long-term in the absence of CD8^+^ T cells, demonstrating that the combined action of innate and adaptive immunity was required for tumour rejection. NK cells play a critical role in the control of orthotopic lung tumours during tumour cell seeding in the lung. To ensure the control of COX-2-deficient tumours was not exacerbated by the route of injection we also compared the growth of subcutaneous parental and *Ptgs2^−/−^* tumours. Similar to the orthotopic setting, COX-2-deficient subcutaneous tumours grew significantly slower in immune-competent mice (Supplementary Fig. 3D). Consistent with the role of both innate and adaptive immunity in the rejection of COX-2-deficient tumours, we observed increased frequencies of CD8^+^ T cells and NK cells as well as CD4^+^ T cells and Tregs in *Ptgs2^−/−^* tumours (Fig. 3C). Increased infiltration of COX-2-deficient tumours by NK cells was confirmed by immunohistochemistry (Fig. 3D).

Further flow cytometry analysis revealed drastic changes in the phenotype of both myeloid and adaptive immune subtypes within COX-2-deficient tumours. T cells infiltrating *Ptgs2*^−/−^ tumours were more activated, with higher frequencies of effector memory (CD44^+^CD62L^−^) CD8^+^ and CD4^+^ T cells (Fig. 3E), mirrored by fewer naïve (CD62L^+^CD44^−^) T cells, an increased frequency of T cells expressing checkpoint receptors (Supplementary Fig. 3E) and upregulation of the activation markers CD44 and CD69 (Fig. 3F, Supplementary Fig. 3F). T cell-mediated immune responses require priming by CD103^+^ dendritic cells (cDC1s) both in lymph nodes and within tumours. Tumour-derived prostaglandins have previously been reported to prevent the accumulation of cDC1s into the TME (33), however we did not see increased cDC1s in *Ptgs2*^−/−^ tumours (Supplementary Fig. 3G). However, we observed increased expression of the co-stimulatory molecule CD86 and MHC-II on cDC1s (Supplementary Fig. 3H). Similarly, loss of COX-2 did not alter the frequency of tumour-associated macrophages (TAMs) (Supplementary Fig. 3I) but induced pro-inflammatory polarisation of both CD11b^+^ and CD11c^+^ TAMs with increased expression of both CD86 and MHC-II (Fig. 3G, Supplementary Fig. 3J) as well as decreased expression of the M2 markers arginase and CD206 (Fig. 3H-I, Supplementary Fig. 3K). Indeed, when assessing the frequency of M1 (MHC-II^+^CD206^−^) and M2 (MHC-II^−^CD206^+^) CD11b^+^ TAMs we observed a dramatic increase of the M1/M2 ratio in COX-2-deficient tumours (Fig. 3J). *Ptgs2*^−/−^ tumours also had fewer immunosuppressive Ly6C^+^ monocytes (Supplementary Fig. 3L). In addition, several myeloid subtypes, including CD11b^+^ TAMs, cDC1s, CDC2s and neutrophils, exhibited increased PD-L1 expression (Supplementary Fig. 3M), characteristic of a T-cell inflamed TME. Gene expression analysis resulted in distinct clustering of parental KPAR and *Ptgs2*^−/−^ tumours with COX-2 loss leading to increased expression of genes encoding Th1 cytokines (*Ifng*, *Tnfa*, *Cxcl9*), cytotoxicity genes (*Gzmb, Prf1*), IFN-response genes (*Irf9, B2m, Stat1*) and decreased expression of the immunosuppressive cytokine IL-6 (Fig. 3K). Together these data suggests that loss of tumour-intrinsic expression of COX-2 results in a remodelling of the tumour microenvironment with increased recruitment of effector cells and polarisation of both innate and adaptive immune subsets towards a pro-inflammatory phenotype resulting in enhanced anti-tumour immunity and greater tumour control.

### COX-2/PGE_2_ signalling drives resistance to ICB in mouse and human LUAD

Given that tumour-intrinsic COX-2 acts as a major driver of immune evasion in KPAR tumours we wanted to assess whether it also contributed to resistance to ICB. Mice were orthotopically transplanted with parental KPAR or *Ptgs2*^−/−^ cells and treated with anti-PD-1. Whilst parental KPAR tumours were partially responsive to anti-PD-1, COX-2 deficient tumours were significantly more sensitive to PD-1 blockade with all ICB-treated mice bearing *Ptgs2*^−/−^ lung tumours surviving long-term (Fig. 4A). Immunohistochemistry revealed that similar to ICB-treated KPAR tumours, *Ptgs2*^−/−^ lung tumours were more infiltrated by CD8^+^ T cells (Fig. 4B), confirming what we observed by flow cytometry (Fig. 3C), however the biggest increase was seen in ICB-treated *Ptgs2*^−/−^ tumours. Furthermore, flow cytometry analysis demonstrated that PD-1 blockade only led to increased activation of CD8^+^ T cells in *Ptgs2*^−/−^ lung tumours (Fig. 4C). In addition, ICB-treated *Ptgs2*^−/−^ lung tumours showed the greatest expansion of effector memory CD8^+^ T cells (Supplementary Fig. 4A) and upregulation of checkpoint molecules (Fig. 4D). Interestingly, anti-PD-1 treatment also led to increased NK cell infiltration in *Ptgs2*^−/−^ lung tumours, which did not occur in KPAR tumours (Supplementary Fig. 4B). Furthermore, gene expression analysis revealed that anti-PD-1 induced robust expression of anti-tumour immunity genes only in COX-2-deficient lung tumours (Fig. 4E, Supplementary Fig. 4C). Together these data suggest that tumour-intrinsic COX-2 promotes resistance to ICB by preventing the stimulation of anti-tumour immunity in response to PD-1 blockade.

**Figure 4.**
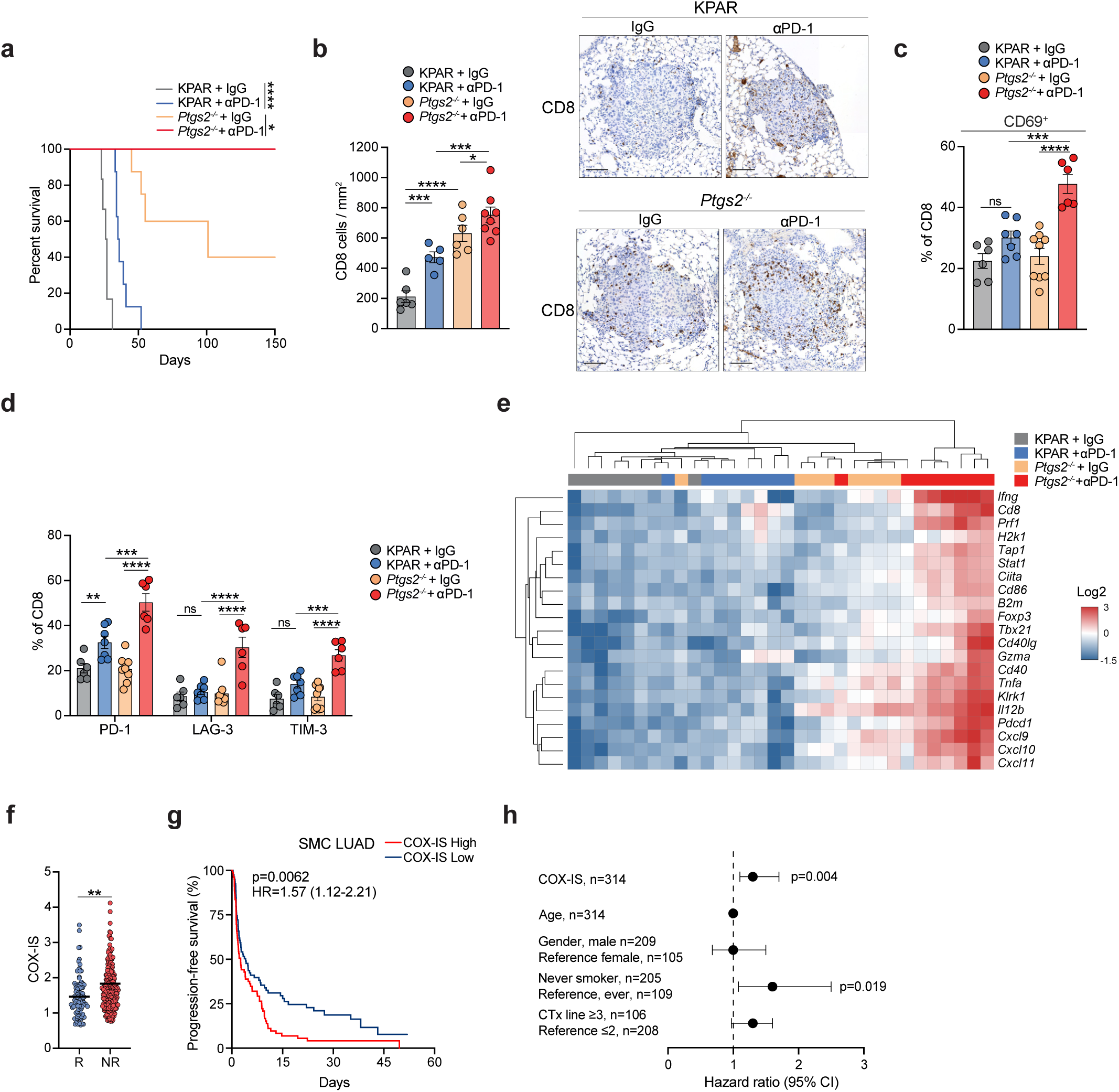
COX-2/PGE2 signalling hinders response to ICB in mouse and human LUAD. (A) Kaplan-Meier survival of mice treated intraperitoneally with 200µg anti-PD-1 after orthotopic transplantation of KPAR or *Ptgs2^−/−^* cells, n=6-8 per group. Analysis of survival curves was carried out using log-rank (Mantel-Cox) test; * P<0.05, **** P<0.0001. (B) Quantification and representative immunohistochemistry staining of CD8+ T cells in KPAR or *Ptgs2^−/−^* orthotopic tumours on day 7 after treatment with anti-PD-1 or corresponding isotype control (IgG). Scale bar represents 100µm. (C) Percentage of CD69^+^ CD8^+^ T cells in KPAR or *Ptgs2^−/−^* tumours treated as in (B) (D) Quantification of PD-1, LAG-3 and TIM-3 surface expression on CD8+ T cells in KPAR or *Ptgs2^−/−^* tumours treated as in (B). (E) Heatmap showing hierarchical clustering of KPAR or *Ptgs2^−/−^* tumours treated as in (B) based on mRNA expression of anti-tumour immunity genes assessed by qPCR. (F) Baseline COX-IS levels in responder and non-responder ICB-treated LUAD patients. (G) Progression-free survival of LUAD patients treated with ICB, stratified into highest and lowest quartile based on COX-IS expression. (H) Multivariate Cox regression analysis for the indicated variables in LUAD patients following ICB treatment (CTx, chemotherapy). Error bars represent 95% confidence interval boundaries. Data are mean ± SEM for (B-D), n=5-9 per group. Statistics were calculated using one-way ANOVA, FDR 0.05; ns, not significant, * P<0.05, ** P<0.01, *** P<0.001, **** P<0.0001.

To determine whether COX-2/PGE_2_ signalling also affected the clinical response of lung cancer patients to immunotherapy we examined the expression of a previously published COX-2-associated inflammatory gene expression signature (COX-IS) (34) in a cohort of LUAD patients treated with anti-PD-L1/PD-1 for which baseline expression data was available (35). Importantly, expression of the COX-IS was significantly higher in LUAD patients who did not respond to ICB (Fig. 4F). Furthermore, higher COX-IS expression was associated with significantly worse progression-free survival following ICB (Fig. 4G) and was also predictive of outcome independent of age, gender, smoking status and previous lines of therapy (Fig. 4H). These results support the notion, as suggested by the mouse model, that the COX-2/PGE_2_ axis drives immunosuppression and hinders response to ICB in human LUAD.

### Inhibition of the COX-2/PGE_2_ axis delays tumour growth and synergises with ICB

Whilst genetic deletion of tumour-intrinsic COX-2 resulted in a drastic repolarisation of the tumour microenvironment, increased tumour control and sensitisation to ICB, we next sought to assess whether pharmacological blockade of COX-2 could have similar effects. We treated KPAR lung tumour-bearing mice with the COX-2 specific inhibitor celecoxib, which was administered by daily oral gavage. As seen in *Ptgs2*^−/−^ lung tumours, treatment of KPAR tumours with celecoxib resulted in polarisation of TAMs with upregulation of CD86 and MHC-II (Fig. 5A, Supplementary Fig. 5A), decreased expression of arginase (Fig. 5B, Supplementary Fig. 5B) and an increase in the M1/M2 ratio (Fig. 5C), as well as reduced infiltration of Ly6C^+^ monocytes (Supplementary Fig. 5C). Changes in the myeloid compartment were accompanied by increased activation of CD8^+^ T cells (Fig. 5D).

**Figure 5.**
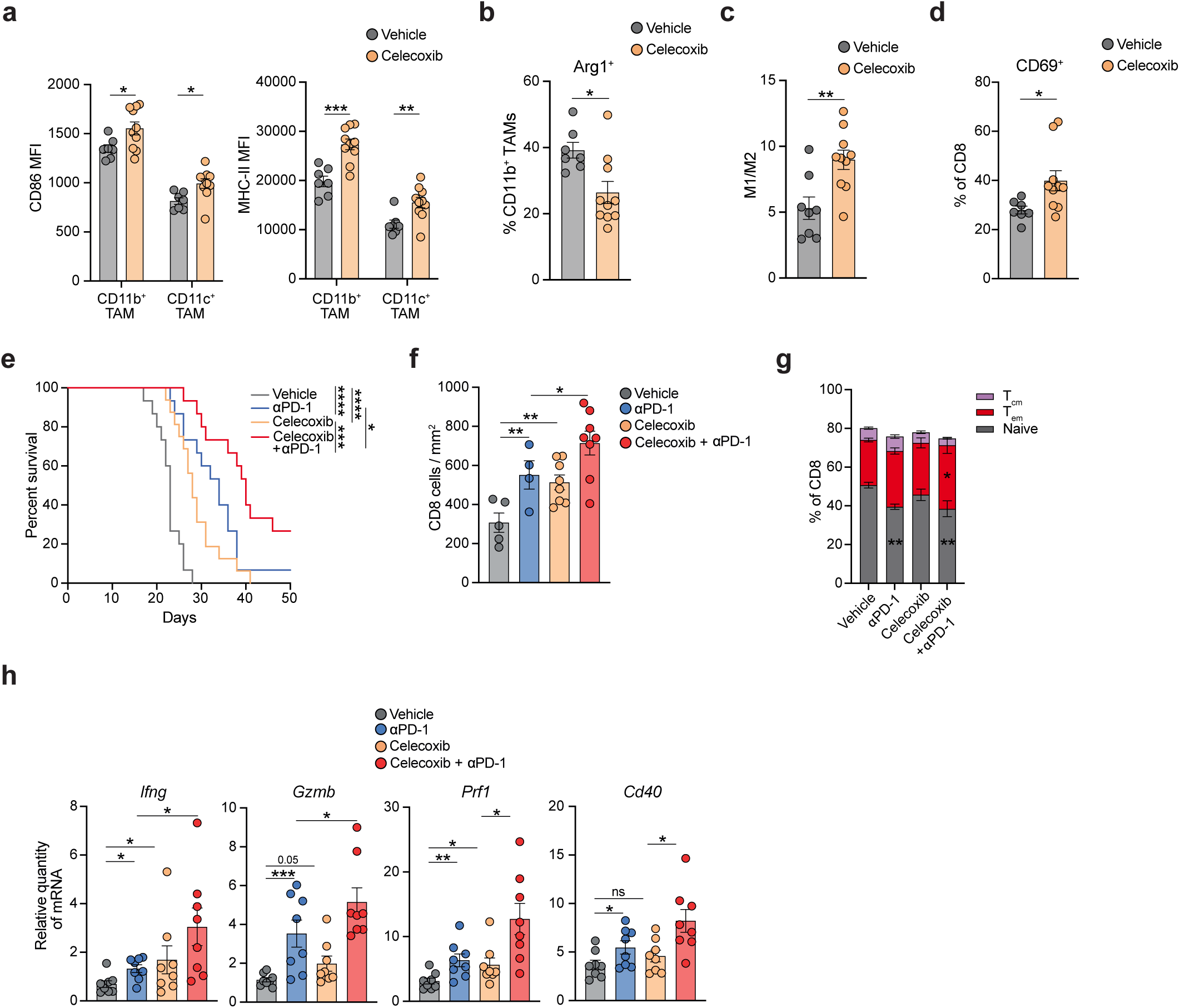
COX-2 inhibition enhances the efficacy of immunotherapy. (A) Surface expression of CD86 (left) and MHC-II (right) on CD11b^+^ macrophages and CD11c^+^ macrophages in KPAR tumours treated for 7d with 30mg/kg celecoxib. (B-D) Percentage of Arg1^+^ CD11b+ macrophages (B), quantification of M1/M2 macrophages (C), and frequency of CD69^+^ CD8^+^ T cells (D) in KPAR tumours treated as in (A). (E) Kaplan-Meier survival of mice treated intraperitoneally with 200µg anti-PD-1 and/or daily oral gavage of 30mg/kg celecoxib after orthotopic transplantation of KPAR cells. Daily celecoxib treatment was initiated on day 7 and anti-PD-1 began on day 10 and was administered twice weekly for a maximum of 3 weeks. Data from two independent experiments, n=15-16 per group. Analysis of survival curves was carried out using log-rank (Mantel-Cox) test; * P<0.05, *** P<0.001, **** P<0.0001. (F) Quantification of CD8+ T cells by immunohistochemistry in KPAR tumours treated for 7d with celecoxib and/or anti-PD-1. (G) CD8^+^ T cell phenotypes in KPAR tumours treated as in (F). (H) mRNA expression by qPCR of anti-tumour immunity genes in KPAR tumours treated as in (F). Data are mean ± SEM for (A-D and F-H), n=5-10 per group. Samples were analysed using unpaired, two-tailed Student’s t-test (A-D), one-way ANOVA, FDR 0.05 (F and H) or two-way ANOVA, FDR 0.05 (G); ns, not significant, * P<0.05, ** P<0.01, *** P<0.001.

Importantly, celecoxib significantly extended the survival of KPAR-tumour bearing mice to a similar extent seen with PD-1 blockade (Fig. 5E). However, the combination of both celecoxib and anti-PD-1 showed superior efficacy compared to either single-agent alone. Indeed, both celecoxib and anti-PD-1 increased infiltration of tumours with CD8^+^ T cells which was further increased in the combination treatment arm (Fig. 5F). Furthermore, only the combination treatment led to a significant expansion of effector memory CD8^+^ T cells (Fig. 5G) and upregulation of checkpoint molecules on both CD8^+^ and CD4^+^ T cells (Supplementary Fig. 5D). Combination treatment also induced the highest levels of PD-L1 on several myeloid cell types (Supplementary Fig. 5E). This could be due to elevated levels of IFNγ (Fig. 5H) which was significantly upregulated in the combination treatment arm along with other anti-tumour immunity genes.

In the clinic, celecoxib has been associated with increased cardiovascular risk (36), which prompted us to explore other therapeutic options to target this immunosuppressive axis. Immune cells express four receptors for PGE_2_, EP1-4. However, numerous studies have demonstrated EP2 and EP4 receptors are the primary mediators of the COX2/PGE_2_ immunosuppressive axis (37,38). We therefore treated tumour-bearing mice with a novel dual EP2-EP4 antagonist TPST-1495 (39). As seen with celecoxib, dual EP2-EP4 inhibition led to a significant increase in the M1/M2 macrophage ratio (Fig. 6A) with reduced expression of arginase (Fig. 6B), as well as increased activation of CD8^+^ T cells (Fig. 6C). Indeed, TPST-1495 increased the survival of tumour-bearing mice similarly to PD-1 blockade (Fig. 6D). The combination of TPST-1495 and anti-PD-1 also significantly extended the survival of mice compared to either monotherapy, with greater synergy as compared to the combination of celecoxib and anti-PD-1. Furthermore, gene expression analysis revealed distinct clustering of tumours treated with the combination with potent induction of a pro-inflammatory transcriptional program (Fig. 6E-F). Consistent with this, the combination treatment led to the biggest increase in PD-L1 expression on tumour-infiltrating myeloid cells (Supplementary Fig. 5F). In conclusion, these results suggest that pharmacological inhibition of the COX-2/PGE_2_ axis reverses immunosuppression in the TME, promoting adaptive immunity which enhances the therapeutic efficacy of ICB.

**Figure 6.**
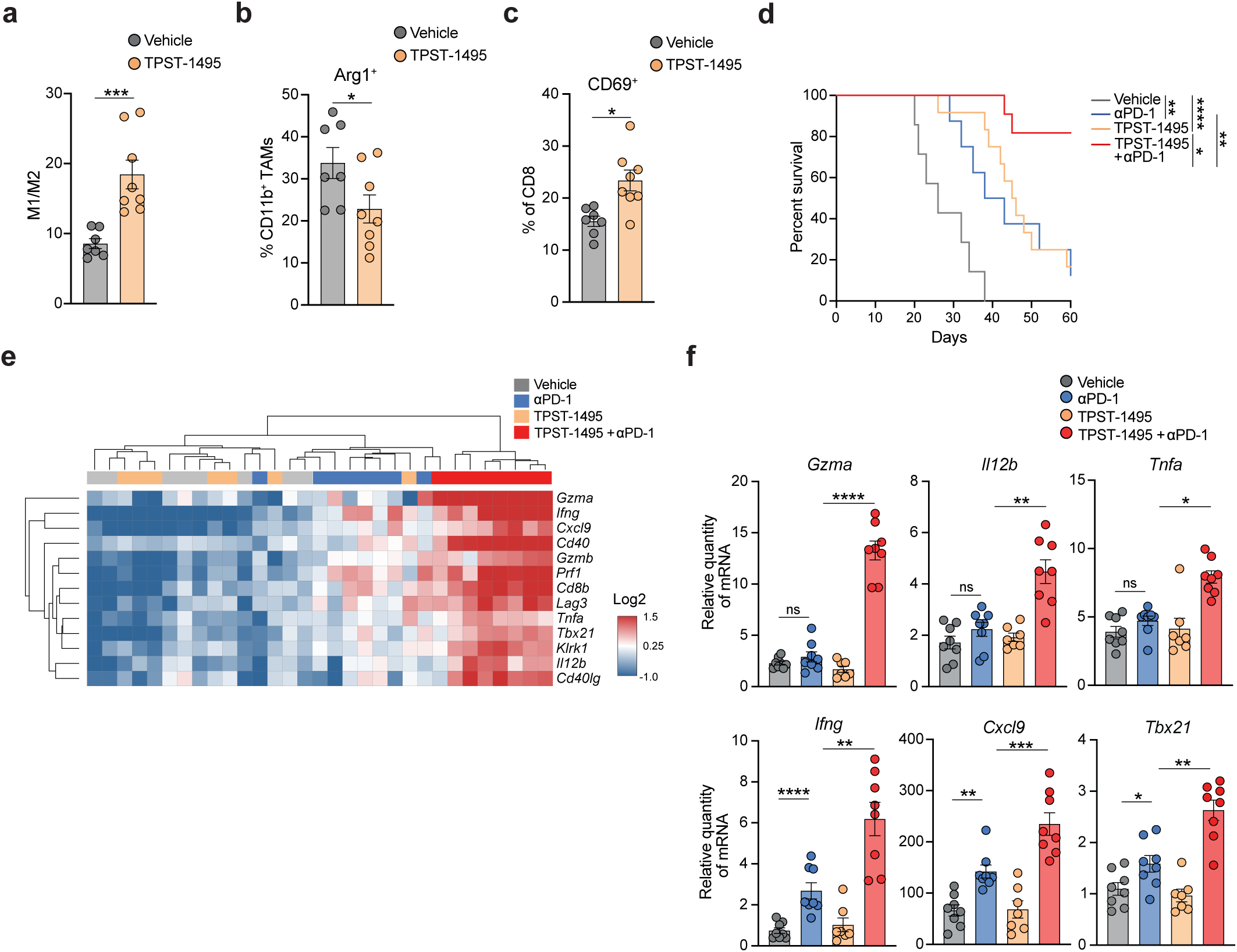
Dual inhibition of EP2 and EP4 synergises with ICB. (A) Kaplan-Meier survival of mice treated intraperitoneally with 200µg anti-PD-1 and/or twice daily oral gavage of 100mg/kg TPST-1495 after orthotopic transplantation of KPAR cells, n=8-12 per group. TPST-1495 treatment was initiated on day 7 and anti-PD-1 began on day 10 and was administered twice weekly for a maximum of 3 weeks. Analysis of survival curves was carried out using log-rank (Mantel-Cox) test; * P<0.05, ** P<0.01, **** P<0.0001. (B) Percentage of CD69^+^ CD8+ (left) and CD4^+^ (right) T cells in KPAR tumours treated for 7d with TPST-1495 and/or anti-PD-1. (C) Heatmap showing hierarchical clustering of KPAR tumours treated as in (B) based on mRNA expression of anti-tumour immunity genes assessed by qPCR. (D) mRNA expression by qPCR of immune-related genes in KPAR tumours treated as in (B). Data are mean ± SEM for (B) and (D), n=7-8 per group. Statistics were calculated using one-way ANOVA, FDR 0.05; ns, not significant, * P<0.05, ** P<0.01, *** P<0.001, **** P<0.0001.

### COX-2 expression is driven by oncogenic KRAS and contributes to tumour relapse after KRAS^G12C^ inhibition

Given the known role of oncogenic KRAS in mediating immune evasion we next wanted to understand whether tumour-intrinsic COX-2 expression was regulated by KRAS signalling. To test this, we inhibited KRAS signalling in several mouse and human KRAS-mutant cancer cell lines. Firstly, treatment of KPAR cells *in vitro* with the MEK inhibitor trametinib led to a drastic reduction in COX-2 protein expression and loss of PGE_2_ secretion (Fig. 7A). We validated this in the 3LL ΔNRAS mouse lung cancer cell line (31), which contains a KRAS^G12C^ mutation and has been rendered sensitive to KRAS^G12C^ inhibitors by deletion of oncogenic NRAS, as well as the CT26^G12C^ colorectal cancer cell line (40) and the KPAR^G12C^ cell line (17) which has been engineered to express KRAS^G12C^. Treatment with the KRAS^G12C^ inhibitor MRTX849 led to reduced COX-2 expression (Fig. 7B) and loss of PGE_2_ secretion in all three cell lines (Fig. 7C). Importantly, celecoxib significantly delayed the growth of 3LL ΔNRAS tumours (Supplementary Fig. 6A) and has recently been shown to reduce the growth of CT26 tumours (38). To extend these findings *in vivo* we carried out gene expression analysis of KP GEMM or KPAR tumours from mice treated with MEKi or TEL, respectively, and observed a decrease in the expression of COX-2 mRNA (Supplementary Fig. 6B-C). MEKi inhibits MAPK signalling in stromal cells as well as tumour cells so we also assessed COX-2 expression levels in 3LL ΔNRAS and KPAR^G12C^ tumours from mice treated with MRTX849, as KRAS^G12C^ inhibitors only target tumour cells. KRAS^G12C^ inhibition downregulated COX-2 mRNA expression in both 3LL ΔNRAS and KPAR^G12C^ tumours (Fig. 7D). This was accompanied by a decrease in COX-2 protein expression (Supplementary Fig. 6D) in KPAR^G12C^ tumours from mice treated with MRTX849. Furthermore, in both tumour models KRAS^G12C^ inhibition reduced the expression of the COX-2-associated inflammatory signature COX-IS (Fig. 7E).

**Figure 7.**
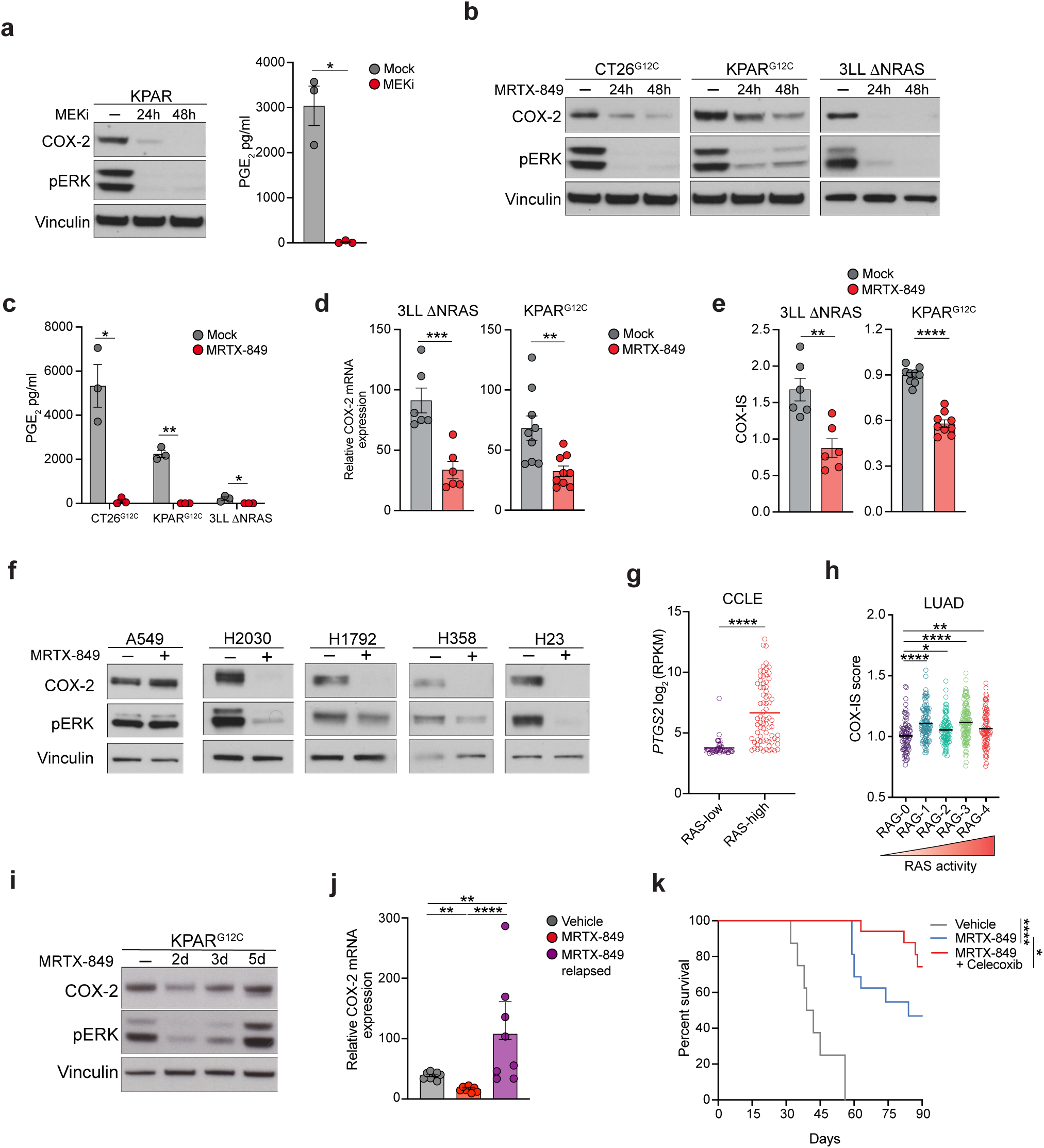
Oncogenic KRAS drives immunosuppressive COX-2 expression in LUAD. (A) Immunoblot for COX-2 (left) and ELISA analysis for PGE2 concentration (right) in KPAR cells treated with 10nM trametinib for 24h or 48h. (B-C) Immunoblot for COX-2 (B) and ELISA analysis for PGE2 concentration (C) in KRAS^G12C^ mouse cancer cell lines treated with 100nM MRTX849 for 24h or 48h. (D) COX-2 mRNA expression in 3LL ΔNRAS and KPAR^G12C^ orthotopic tumours treated for 7d with 50mg/kg MRTX849 (E) COX-2-associated inflammatory signature (COX-IS) assessed by qPCR in 3LL ΔNRAS and KPAR^G12C^ orthotopic tumours treated as in (D). (F) Immunoblot for COX-2 in human KRAS^G12C^ lung cancer cell lines treated with MRTX849 for 24h. A549 (KRAS^G12S^) cells were used as a negative control. (G) COX-2 expression in RAS-low and RAS-high human lung cancer cell lines from the CCLE database. (H) COX-IS in LUAD samples from TCGA stratified by RAS-activity into five different groups which are associated with specific co-occurring mutations (RAG-1, KRAS/LKB1; RAG-2, KRAS; RAG-3, KRAS/TP53; RAG-4, KRAS/CDKN2A). (I) Immunoblot for COX-2 in KPAR^G12C^ cells treated for 2d, 3d or 5d with 100nM MRTX849 (J) COX-2 mRNA expression in MRTX849 on-treatment and relapsed KPAR^G12C^ tumours. (K) Kaplan-Meier survival of mice treated with daily oral gavage of 50mg/kg MRTX849 alone or in combination with 30mg/kg celecoxib, n=8-20 per group. Analysis of survival curves was carried out using log-rank (Mantel-Cox) test; * P<0.05, **** P<0.0001. Data are mean ± SEM for (A, C-E and J), n=8-9 per group. Groups were compared using unpaired, two-tailed Student’s t-test (A, C-E and G) or one-way ANOVA, FDR 0.05 (H and J); * P<0.05, ** P<0.01, ***, P<0.001, **** P<0.0001.

Consistent with our results in mouse cancer cell lines, we also observed significant downregulation of COX-2 expression in a panel of KRAS^G12C^ human LUAD cell lines treated *in vitro* with MRTX849 (Fig. 7F). Such an effect was not observed in KRAS^G12S^ A549 cells which are insensitive to KRAS^G12C^ inhibition. We extended our analysis using expression data from human lung cancer cell lines from the Broad Institute Cancer Cell Line Encyclopedia (CCLE) which were classified as either having high or low oncogenic RAS pathway activity based on a recently published RAS transcriptional signature (26). We observed significantly increased expression of *PTGS2* in high-RAS pathway activity cell lines (Fig. 7G). Similarly, *PTGS2* expression was increased in KRAS-mutant cell lines (Supplementary Fig. 6E). We also analysed LUAD expression data from TCGA which were classified into five different groups, RAG-0 to RAG-4, based on expression of the RAS activity transcriptional signature, with RAG-0 being the lowest and RAG-4 being the highest. COX-IS was significantly increased in all RAS active groups with highest levels occurring in RAG-1 and RAG-3 which are associated with co-occurring *STK11/LKB1* or *TP53* mutations, respectively (Fig. 7H). Similarly, *PTGS2* mRNA levels were elevated in RAS active tumours as well as KRAS-mutant tumours (Supplementary Fig. 6F). Interestingly, COX-IS was not elevated in KRAS-mutant tumours (Supplementary Fig. 6G) demonstrating the advantage of utilising a RAS transcriptional signature, rather than simply KRAS mutation status, to capture RAS signalling activity in human lung cancer.

The advent of KRAS^G12C^ inhibitors has transformed the treatment landscape for KRAS^G12C^-mutant lung cancer patients. However, responses are often short-lived and combination therapies will be required to overcome the development of adaptive resistance. Whilst oncogenic KRAS pathway reactivation will re-engage proliferative signalling we postulated that restoration of KRAS-mediated immunosuppression may also contribute to tumour relapse. Interestingly, COX-2 expression was restored in KPAR^G12C^ cells *in vitro* after long-term MRTX849 treatment (Fig. 7I) and *in vivo* in KPAR^G12C^ tumours that relapsed after MRTX849 treatment (Fig. 7J). Importantly, the combination of MRTX849 and celecoxib delayed tumour relapse and significantly improved the survival of KPAR^G12C^ tumour-bearing mice compared to MRTX849 alone (Fig. 7K).

In summary, these data suggest that oncogenic KRAS is a major driver of COX-2/PGE_2_ signalling in mouse and human lung cancer and may therefore contribute to ICB resistance in KRAS-mutant LUAD and tumour relapse in patients treated with KRAS^G12C^ inhibitors which can be overcome by COX-2 pathway inhibitors.

## DISCUSSION

A subset of KRAS-mutant lung cancer patients has greatly benefited from the advent of immune checkpoint blockade therapy. However, the majority of patients still do not respond. Combination strategies are urgently required broaden the efficacy of immunotherapy. Accumulating evidence suggest tumour-intrinsic oncogenic signalling can dampen anti-tumour immune responses (26) which could be therapeutically exploited to overcome immunotherapy resistance in specific tumour subtypes, but little progress has been made to date in attempts to combine immunotherapies with therapies targeting oncogenic signalling pathways. Here, we carried out *in vivo* CRISPR screening using a novel immunogenic model of KRAS-mutant LUAD (17) to uncover multiple mechanisms by which oncogenic KRAS drives immune evasion, with a view to inform the development of optimal combination strategies for targeting KRAS mutant cancers that impact both oncogenic signalling and immune evasion.

We demonstrated that tumour-intrinsic IFNγ signalling is critical for anti-tumour immunity. Indeed, numerous CRISPR screens have demonstrated that defects in IFNγ signalling result in resistance to immunotherapy (21,41) and mutations in *JAK1* and *JAK2* have been associated with acquired resistance to ICB (42). Given the importance of tumour-intrinsic IFNγ signalling in sensitising KPAR tumours to anti-tumour immune responses, the ability of oncogenic KRAS to suppress IFN pathway signalling may represent a major mechanism of immune evasion in KRAS-mutant LUAD. These results are consistent with our previous findings that KRAS^G12C^ inhibitors can restore tumour-intrinsic IFN signalling in multiple preclinical models of lung cancer (15). Paradoxically, tumour-intrinsic IFNγ signalling has also been shown to impede anti-tumour immunity (43), including in a CRISPR screen using an orthotopic KRAS^G12D^ p53^−/−^ lung cancer model similar to the one used in this study (25). Whilst such opposing effects in similar models are surprising, chronic tumour-intrinsic IFNγ signalling can drive resistance to ICB due to upregulation of immune checkpoint ligands (44). Understanding the contexts in which tumour-intrinsic IFNγ signalling promotes or impedes anti-tumour immunity will be important when assessing combination strategies targeting this pathway.

We also identified KRAS-driven expression of COX-2 and secretion of PGE_2_ as a major mechanism of immune evasion which suppresses both innate and adaptive anti-tumour immune responses. This is consistent with the role of tumour-intrinsic COX-2 in driving immune evasion in melanoma and colorectal cancer (33). The ability of our CRISPR screen to elucidate the function of a secreted molecule can be explained by a recent barcoded CRISPR screen which demonstrated that orthotopic KP tumours grow as clonal lesions (25), suggesting that local secretion of a particular molecule by neighbouring tumour cells that do not contain the same gene deletion is unlikely to be a major problem.

COX-2 derived PGE_2_ is a pleiotropic molecule that has been shown to act on many cell types including CD8^+^ T cells, dendritic cells and NK cells (45) although single-cell sequencing studies have revealed EP2 and EP4 receptors to be highly expressed specifically on tumour-infiltrating myeloid cells (46). Indeed, we observed extensive changes in the TME of COX-2-deficient tumours including pro-inflammatory polarisation of myeloid cells and concomitant infiltration of highly activated CD8^+^ T cells and NK cells. Interestingly, NK cell depletion was best able to restore the growth of COX-2-deficient tumours, suggesting that NK cells may play a major role in the rejection of COX-2-deficient tumours. This supports observations that NK cells drive remodelling of the tumour microenvironment and promote immune-control in COX-2-deficient melanoma tumours (34). Furthermore, the effects we observed of tumour-derived PGE_2_ on myeloid cells is consistent with a recent study that demonstrated the immunosuppressive role of PGE_2_-producing lung fibroblasts which promote breast cancer metastasis (37). The role of COX-2 expressing lung fibroblasts within primary lung tumours remains unclear, however our results suggest that the majority of PGE_2_ secretion within the TME is derived from tumour cells as tumour-specific genetic deletion of COX-2 was sufficient to drive immune-mediated tumour eradication. Unlike studies in melanoma (33), we did not observe increased cDC1 recruitment in COX-2-deficient tumours, reflecting the context-specific effects of prostaglandin signalling in different tumour types. cDC1s have a unique ability to prime CD8^+^ T cells and the increased expression of co-stimulatory molecule expression on cDC1s in *Ptgs2*^−/−^ tumours may explain the enhanced T cell activation observed. A recent study utilising the syngeneic 3LL/LLC lung cancer model described a role for Tregs in mediating the immunosuppressive effects of the COX-2/PGE_2_ signalling axis in subcutaneous tumours (46) which we did not observe in our experimental model system. This may be explained by differences in anti-tumour immune responses observed in orthotopic versus subcutaneous tumours. Indeed, multiple transplantable lung cancer models have been shown to have abundant infiltration of Tregs in subcutaneous tumours compared to orthotopic lung tumours (17,47).

Our data demonstrates the therapeutic benefits of genetic ablation or pharmacological inhibition of COX-2 in a preclinical model of KRAS-mutant lung cancer, especially in combination with immunotherapy. Furthermore, the ability of the COX-IS to independently predict outcome after ICB suggests a role for COX-2/PGE_2_ signalling in hindering responses to ICB in LUAD patients, supporting previous findings in other cancer types (34). Given that the expression of the COX-IS was strongly driven by oncogenic KRAS signalling in both mouse and human lung cancer, our studies suggest that inhibiting the COX-2/PGE_2_ axis is a promising therapeutic strategy in KRAS-mutant NSCLC and may broaden the efficacy of immune checkpoint blockade for these patients. Interestingly, our analysis of COX-IS expression in the LUAD TCGA cohort demonstrated increased activity of the COX-2/PGE_2_ axis in RAG-1 patients, which are associated with STK11/LKB1 co-occurring mutations. These patients respond poorly to ICB (48) and therefore may particularly benefit from this combination therapy.

Indeed, a number of clinical trials are currently testing the combination of celecoxib with immune checkpoint blockade (NCT03026140, NCT03864575). However, these trials do not specifically enrol KRAS-mutant LUAD patients which our data suggests would most benefit from such a combination. The potential benefit of this combination is supported by a recent retrospective analysis showing improved survival of lung cancer patients that were concurrently treated with COX inhibitors whilst receiving immunotherapy (49). Furthermore, we show using a novel drug that dual inhibition of the PGE_2_ receptors EP2 and EP4 has similar therapeutic benefits to COX-2 inhibition and shows superior synergy when combined with ICB. This supports other work in preclinical models that have demonstrated the benefits of combining EP2 and EP4 antagonists with ICB (37,38). EP2-4 inhibition therefore has the potential to enhance the efficacy of immunotherapy in the clinic, whilst possibly avoiding the toxicities associated with celecoxib treatment.

The ability of KRAS^G12C^ inhibition to suppress the COX-2/PGE_2_ signalling axis *in vivo* may in part explain the synergy observed in combination with ICB in preclinical models (15,17,18,40). However, given the apparent poor tolerability of the combination of KRAS^G12C^ inhibition by sotorasib and PD(L)-1-targeted ICB observed in the clinic (50) it may be more feasible to target KRAS-driven immune evasion mechanisms such as COX-2. The benefits of KRAS^G12C^ inhibition in the clinic are also seriously confounded by the rapid emergence of acquired resistance which can be driven by many different oncogenic mutations within the RAS signalling pathway (11). Clinical trials such as CODEBREAK 101 are attempting to overcome this by combining KRAS^G12C^ inhibitors with other targeted therapies such as RTK inhibitors; however, given the genetic complexity underlying resistance, the feasibility of this approach remains unclear. Instead, targeting KRAS-driven immune suppression may prove more successful. Our work shows that long-term KRAS^G12C^ inhibition results in restoration of the COX-2/PGE_2_ axis which may contribute to tumour relapse. Furthermore, our data suggest that combination of KRAS^G12C^ inhibitory drugs and COX-2 or EP2-4 prostaglandin receptor inhibition may be successful in the treatment of immune hot lung cancer. One might speculate that it could possibly avoid the toxicities reported for sotorasib and PD(L)-1 blockade.

## METHODS

### *In vivo* tumour studies

All animal studies were approved by the ethics committee of the Francis Crick Institute and conducted according to local guidelines and UK Home Office regulations under the Animals Scientific Procedures Act 1986 (ASPA). All transplantation animal experiments were carried out using 8-12-week C57BL/6J mice. For orthotopic experiments, mice were injected intravenously into the tail-vein with 1.5×10^5^ KPAR or KPAR^G12C^ cells. Mice were euthanised when mice displayed signs of ill health or exceeded 15% weight loss. For subcutaneous experiments, 1×10^6^ 3LL ΔNRAS cells were injected subcutaneously into the left flank (1:1 mix with Matrigel). Tumours were measured twice weekly using callipers and volume calculated using the formula 0.5 × length × width^2^. Mice were euthanised when the average tumour dimension exceeded 1.5 cm.

For antibody treatments, 200μg anti-PD-1 (clone RMP1-14, BioXcell) or the respective IgG control, were administered via intraperitoneal injection twice weekly for a maximum of three weeks. For drug treatments, 50 mg/kg MRTX849 (MedChem Express), 1.3 mg/kg trametinib (LC laboratories), 16.6 mg/kg linsitinib (Astellas), 1.6 mg/kg everolimus (LC laboratories), 30 mg/kg celecoxib (LC laboratories), 100 mg/kg TPST-1495 (kindly provided by Tempest Therapeutics) or their respective vehicles were administered daily or twice daily via oral gavage for the stated duration. MRTX849 was prepared in 10% Captisol diluted in 50mM citrate buffer (pH 5.0), trametinib, everolimus and linsitinib in 0.5% methylcellulose/0.2% Tween 80, celecoxib in a mixture of 10% DMSO, 50% polyethylene glycol 400, and 40% water and TPST-1495 in 0.5% methylcellulose.

For depletion experiments, mice received 200µg anti-CD8 (2.43) and/or 200µg anti-NK1.1 (PK136) via intraperitoneal injection one and three days before tumour cell transplantation followed by once weekly for the duration of the experiment. Depletion was confirmed by flow cytometry using anti-CD49b-AF488 (DX5, Biolegend) and anti-Nkp46-BV421 (29A1.4, Biolegend) for NK cells and anti-CD8-PE (53-6.7, BD Biosciences) for CD8^+^ T cells.

### Cell lines

KPAR, KPAR^G12C^, 3LL ΔNRAS were generated as previously described (17,31). CT26^G12C^ were kindly provided by Mirati Therapeutics (40). NCI-H23, NCI-H358, NCI-H1792, NCI-H2030 and A549 were obtained from the Francis Crick Institute Cell Services Facility. Cell lines were cultured in DMEM or RPMI supplemented with fetal bovine serum (10%), L-glutamine (2mM), penicillin (100 units/mL) and streptomycin (100 µg/mL). Clonal cell lines were derived by single-cell dilution into 96 well plates in DMEM-F12 supplemented with GlutaMAX, FBS (10%), hydrocortisone (1µM), EGF (20 ng/ml) and IGF (50ng/ml). Cell lines were routinely tested for mycoplasma infection and were authenticated by short-tandem repeat (STR) DNA profiling by the Francis Crick Institute Cell Services facility. For *in vitro* growth experiments, 2×10^4^ cells were plated in 96-well plates and growth was monitored for 5d using an IncuCyte Zoom system (Essen Biosciences).

### In vitro treatments

Drugs or cytokines were added in fresh media 24h after seeding cells at stated concentrations and samples were obtained at indicated time points.

### Flow cytometry

Mouse tumours were cut into small pieces and incubated with collagenase (1 mg/ml; ThermoFisher) and DNase I (50 U/ml; Life Technologies) in HBSS for 45 min at 37°C. Cells were filtered through 70 μm strainers (Falcon) and red blood cells were lysed using ACK buffer (Life Technologies). Samples were stained with fixable viability dye eFluor870 (BD Horizon) for 30 min and blocked with CD16/32 antibody (Biolegend) for 10 min before fluorescently-labelled antibody staining of surface markers (see Supplementary Table 2, Supplementary Fig. 7). Intracellular staining was then performed after fixation using the Fixation/Permeabilization kit (eBioscience) according to the manufacturer’s instructions. Samples were resuspended in FACS buffer and analysed using a BD Symphony flow cytometer. Data was analysed using FlowJo (Tree Star).

For FACS analysis *in vitro*, cells were trypsinised, washed with FACS buffer and stained with the following antibodies: anti-IFNγR-β-chain-PE (MOB-47), anti-H2Db-PE (KH95) or anti-PD-L1-PE-Cy7 (10F.9G2).

### Immunohistochemistry

Tumour-bearing lungs were fixed in 10% NBF for 24h and transferred to 70% ethanol. Fixed lungs were processed into paraffin-embedded blocks. Tissue sections were stained with haematoxylin and eosin, using the automated TissueTek Prisma slide stainer. For immunohistochemistry staining, sections were boiled in sodium citrate buffer (pH 6.0) for 15 min and incubated with the following antibodies for 1h: anti-CD8 (4SM15, Thermo Scientific), anti-NCR1 (EPR23097-35, Abcam) and anti-Arg1 (D4E3M, Cell Signalling). Primary antibodies were detected using biotinylated secondary antibodies and HRP/DAB detection. Slides were imaged using a Leica Zeiss AxioScan.Z1 slide scanner. Tumour-infiltrating immune cells were quantified using QuPath.

### RT-qPCR

RNA was extracted from cell lines or frozen lung tumours using RNeasy kit (Qiagen). Single tumour nodules were plucked from tumour-bearing lungs. Tumours from multiple mice were included in each analysis. Frozen tumour samples were homogenised prior to RNA extraction using QIAshredder columns (Qiagen). cDNA was generated using the Maxima First Strand cDNA Synthesis Kit (Thermo Fisher Scientific) and qPCR performed using Applied Biosystems Fast SYBR Green reagents (Thermo Fisher Scientific). mRNA relative quantity was calculated using the ΔΔCT method and normalised to *Sdha*, *Hsp90* and *Tbp*. For heatmap visualisation, relative mRNA expression for each gene was log-transformed and median-normalised.

### ELISA

Cells were treated as indicated for 48h. Conditioned media was collected and PGE_2_ concentration determined using the Prostaglandin E2 parameter assay kit (R&D), as per manufacturer’s instructions.

### siRNA experiments

Cells were seeded in a 6-well plate and reverse-transfected with 50nM siGENOME siRNA pools targeting mouse *Myc* (Dharmacon) using DharmaFECT 4 transfection reagent (Dharmacon) according to the manufacturer’s instructions. 24h after transfection, cells were treated for 24h with trametinib (10nM). Control cells were mock-transfected (no siRNA) or transfected with siGENOME RISC-free control siRNA (Dharmacon).

### MicroCT imaging

Mice were anesthetised by inhalation of isoflurane and scanned using the Quantum GX2 micro-CT imaging system (Perkin Elmer). Serial lung images were reconstructed and tumour volumes subsequently analysed using Analyse (AnalyzeDirect) as previously described (51)

### Western blotting

Cells were lysed using 10X Cell Lysis Buffer (Cell Signalling) supplemented with protease and phosphatase inhibitors (Roche). Protein concentration was determined using a BCA protein assay kit (Pierce) and 15-20 μg of protein was separated on a 4-12% NuPAGE Bis-Tris gel (Life Technologies) followed by transfer to PVDF membranes. Proteins were detected by Western blotting using the following primary antibodies against: Flag (M2, Sigma), ERK1/2 (3A7, Cell Signalling), p-ERK1/2 (Thr202/Tyr204) (9101, Cell Signalling), Myc (Y69, Abcam), STAT1 (#9172, Cell Signaling), p-STAT1 (T701) (58D6, Cell Signaling) STAT2 (D9J7L, Cell Signaling) COX-2 (D5H5, Cell Signalling), Vinculin (VIN-11-5, Sigma) and β-Actin (8H10D10, Cell Signaling). Primary antibodies were detected using HRP-conjugated secondary antibodies and visualised using standard film.

### CRISPR/Cas9 knockout

*Ptgs2*^−/−^, *Ifngr2*^−/−^ and *Etv4*^−/−^ KPAR cell lines were generated by transient transfection of a Cas9-sgRNA plasmid (pX459, Addgene) generated by standard molecular cloning techniques (see Supplementary Table 3). 3×10^5^ cells were seeded in a 6-well plate and transfected 24h later with 5 µg pX459 plasmid DNA using Lipofectamine 3000 (Thermo Scientific) according to the manufacturer’s instructions. After 24h, cells were selected in puromycin (5 µg/ml, InvivoGen) for 48h. After selection, cells were single-cell cloned by single-cell dilution into 96 well plates. Knockout clones were identified by Western blot, ELISA or flow cytometry. *Edn1*^−/−^ KPAR cells were generated by infecting KPAR iCas9 cells with a pool of five different lentivirus each encoding a different sgRNA cloned into a lentiviral vector (pLenti_BSD_sgRNA). Cells were infected at a high multiplicity of infection (MOI) to maximise the number of lentivirus particles taken up by cells and increase the efficacy of editing. 24h after infection, cells were selected in blasticidin (10 µg/ml, InvivoGen) for 4 days and Cas9 induced by doxycycline (1 µg/ml, Sigma) for 6 days. As a negative control, cells were infected with a pool of five different lentivirus each encoding a different non-targeting sgRNA. sgRNAs were designed using the GPP sgRNA designer (http://portals.broadinstitute.org/gpp/public/ analysis-tools/sgrna-design).

### Stable cell lines, plasmids and lentivirus infection

For generation of the KPAR iCas9 cell line, KPAR cells were infected with the lentiviral vector pCW-Cas9 (Addgene) at a MOI of 0.3. 24h after infection, cells were selected in hygromycin (500 µg/ml, InvivoGen) for 7 days. Antibiotic-selected cells were single-cell cloned by single-cell dilution in 96 well plates. Clones with minimal expression of Cas9 in normal media and robust induction of Cas9 expression after 24h treatment with doxycycline (1 µg/ml, Sigma) were identified by Western blotting. Lentivirus particles were generated by co-transfection of HEK293T cells with the lentiviral vector and packaging plasmids pCMV-VSV-G and pCMV-dR8.2. 48h after transfection, supernatant was collected, filtered through a 0.45µm filter and frozen at −80°C. Cells were infected with lentivirus particles by spinfection. Briefly, 1×10^6^ cells were plated in a 12-well plate along with 8 µg/ml polybrene (Millipore) and the specific volume of lentivirus depending on the MOI desired. Cells were centrifuged at 1000*g* for 2h at 33°C. 2ml of media was added after the spin and 16h later cells were trypsinised and plated into 6-well plates. 24h after spinfection cells were selected with appropriate antibiotic and subsequently expanded.

MOI was calculated for each lentivirus batch by infecting target cells with different dilutions of lentivirus as previously described (52). 16h after spinfection cells were trypsinised and 5×10^4^ cells were plated into 6-well plates followed by selection with the appropriate antibiotic for 4-5 days. Cells were subsequently counted and percent transduction was calculated for each virus dilution by dividing the cell count of antibiotic-selected cells with the cell count of a no-selection control.

### sgRNA library generation

sgRNAs were designed to target upstream of the first functional domain of each gene. This was identified for each gene using the CDS of the principal isoform (APPRIS database) for each transcript and protein domain annotation from the Pfam database. The top five ranked sgRNAs, based on on-target and off-target activity, were chosen. Some genes could only be targeted by four sgRNAs due to their short length. 1,191 sgRNAs, targeting 240 genes, were designed using the GPP sgRNA designer (http://portals.broadinstitute.org/gpp/public/analysis-tools/sgrna-design). sgRNA sequences along with on-target and off-target scores are stated in Appendix 1. 50 non-targeting sgRNAs were also used from the mouse GeCKOv2 library. Annealed oligonucleotides corresponding to each sgRNA were combined into 10 different pools, each containing 125 sgRNAs and 5 non-targeting controls, and cloned into a lentiviral vector (pLenti_BSD_sgRNA). Lentiviral libraries were produced using each of the 10 pools and stored at −80°C.

### *In vivo* CRISPR screening

1×10^6^ KPAR iCas9 cells were infected with each of the 10 lentivirus pools at an MOI of 0.3. 24h after infection, cells were selected in blasticidin (10µg/ml, InvivoGen) for 4 days. Selected were subsequently expanded *in vitro* and Cas9 induced by doxycycline (1 µg/ml, Sigma) for 6 days. Cells were then changed into normal media for 2 days before being orthotopically transplanted into mice. 1.25×10^5^ library-transduced cells were injected intravenously into the tail-vein of wild-type C57BL/6J mice and *Rag2^−/−^*; *Il2rg^−/−^* mice. In parallel, library-transduced cells were cultured *in vitro* at a library representation of >2000x for the same time period as the *in vivo* experiment. Tumours were harvested 2-3 weeks after transplantation and genomic DNA was extracted from tumours using the Gentra Puregene DNA Extraction kit (QIAGEN).

Two-step PCR of genomic DNA was performed to amplify the sgRNA sequences and attach sequencing adaptors required for Illumina sequencing. sgRNA representation in each sample was measured by sequencing amplicons using an Illumina NextSeq 500. Data analysis was performed by the bioinformatics facility at the Francis Crick Institute. Briefly, reads were initially assigned to each sample using the indexed barcodes and then aligned to one of the possible sgRNAs in the library. Reads were normalised to total read counts per sample using MAGeCK and log_2_-fold change between groups calculated.

### Bioinformatic analysis

The CCLE RNA-seq data were obtained from the CCLE repository hosted at The Broad (https://data.broadinstitute.org/ccle_legacy_data). We used the classification of RAS-low and RAS-high as previously described (26). All TCGA RNA-Seq gene-level read counts were downloaded using the TCGAbiolinks (TCGAbiolinks_2.8.4) package from Bioconductor (legacy=TRUE). The samples we classified in RAGs using RAS84, as previously shown (26). Raw counts for the TCGA LUAD, CCLE lung cell lines and LUAD ICB cohort were VST normalised using the varianceStabilizingTransformation function within DESeq2 (DESeq2_1.20.0) from Bioconductor.

The COX-IS score was calculated as the mean expression (vst estimate) of the COX-IS cancer-promoting genes (*VEGFA, CCL2, IL8, CXCL1, CXCL2, CSF3, IL6, IL1B* and *IL1A*) divided by the mean expression (vst estimate) of the COX-IS cancer-inhibitory genes (*CCL5, CXCL9, CXCL10, CXCL11, IL12A, IL12B, IFNG, CD8A, CD8B, GZMA, GZMB, EOMES, PRF1, STAT1* and *TBX21*), as previously described (34). The COX-IS was used to stratify ICB-treated LUAD patients into top 25% and bottom 25% quartiles and univariate survival analysis carried out. Responders were defined as patients with partial or complete response and non-responders were defined as patients with stable or progressive disease.

### Statistical analysis

Statistical significance was assessed in Prism 7 (GraphPad Software) using either an unpaired, two-tailed Student’s t-test, log-rank test, one-way ANOVA or two-way ANOVA, as indicated. P≤0.05 were considered statistically significant (* P<0.05, ** P<0.01, ***P<0.001, **** P<0.0001).

## ACKNOWLEDGEMENTS

We thank the science technology platforms at the Francis Crick Institute including Biological Resources, Advanced Sequencing, Scientific Computing, Bioinformatics and Biostatistics, Flow Cytometry, Experimental Histopathology, and Cell Services.

## Funding

This work was supported by the Francis Crick Institute which receives its core funding from Cancer Research UK (FC001070), the UK Medical Research Council (FC001070), and the Wellcome Trust (FC001070). This work also received funding from the European Research Council Advanced Grant RASImmune, and from a Wellcome Trust Senior Investigator Award 103799/Z/14/Z.

## Competing interests

J.D. has acted as a consultant for AstraZeneca, Jubilant, Theras, Roche and Vividion and has funded research agreements with Bristol Myers Squibb, Revolution Medicines and AstraZeneca. S.C.T has acted as a consultant for Revolution Medicines. K.L. has a patent on indel burden and CPI response pending and speaker fees from Roche tissue diagnostics, research funding from CRUK TDL/Ono/LifeArc alliance, and a consulting role with Monopteros Therapeutics. The other authors declare that they have no competing interests.

## Author contributions

J.B, M.M-A and J.D. designed the study, interpreted the results and wrote the manuscript. J.B., A.D-C, S.R, P.A and E.M performed the biochemical experiments, C.M. assisted with *in vivo* studies, N.B, H.C, S.C.T, P.E and R.G. performed bioinformatics analyses, S-H.L and K.L provided patient data. All authors contributed to manuscript revision and review.

## Notes

### Summary of Updates

Minor revisions have been made throughout to improve data quality and presentation.

