## Supplementary Figures 1-7 for "CRISPR/Cas9 screen identifies KRAS-induced COX-2 as a driver of immunotherapy resistance in lung cancer"

### Supp Figure 1

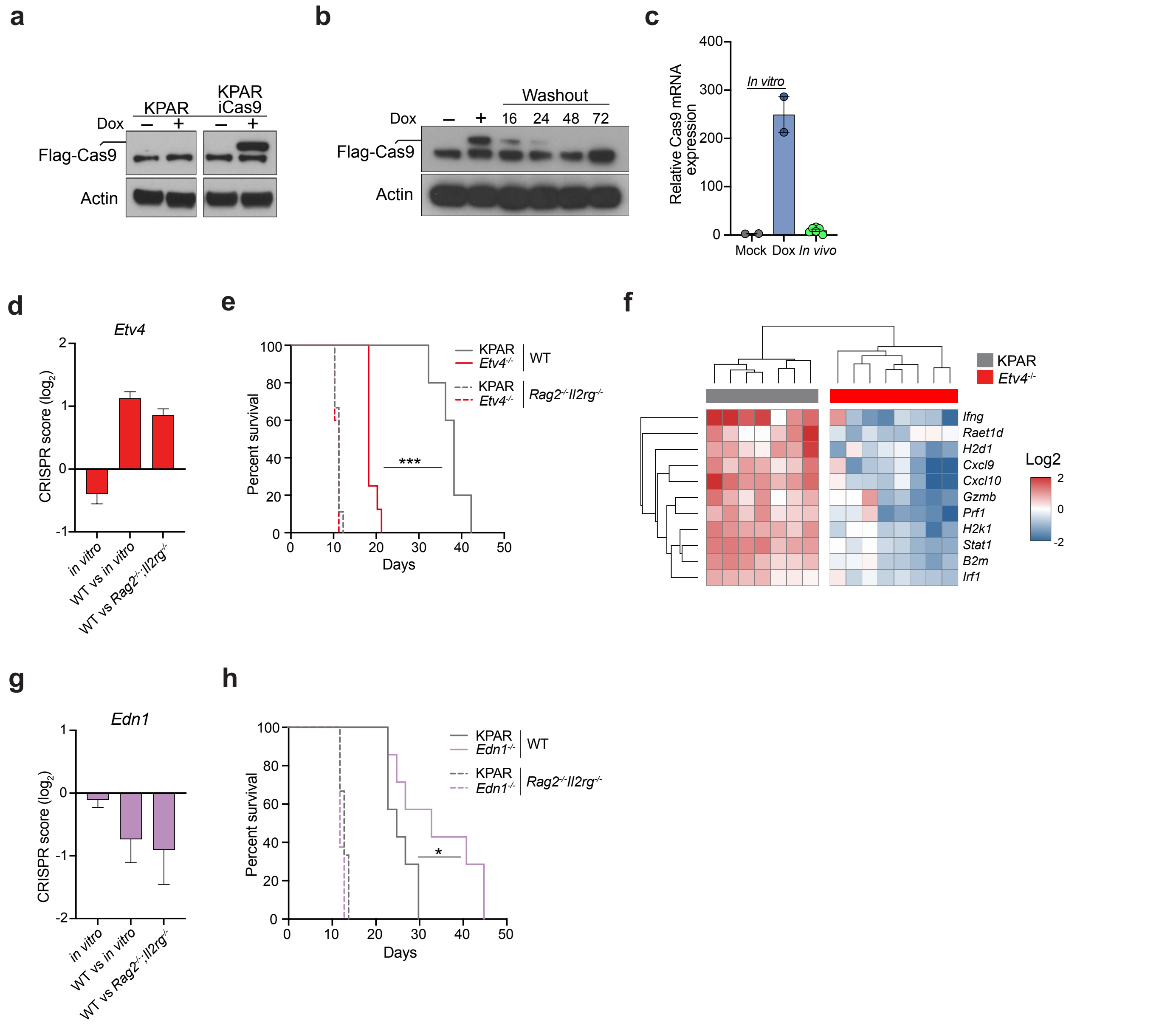


**Supplementary Figure 1. In vivo screen identifies mediators of immune resistance and sensitivity**
(A) Immunoblot for Flag-Cas9 in KPAR iCas9 cells treated for 24h with 1μM doxycycline. Parental KPAR cells were used as a control.
(B) Immunoblot for Flag-Cas9 in KPAR iCas9 cells treated as in (A) followed by washout at stated time points.
(C) mRNA expression by qPCR of Cas9 in KPAR iCas9 cells treated in vitro with 1μM doxycycline or KPAR iCas9 subcutaneous tumours.
(D) Enrichment of sgRNAs targeting Etv4 in WT versus *Rag2*$^{-/-}$*;Il2rg*$^{-/-}$ mice.
(E) Kaplan-Meier survival of immune-competent or *Rag2*$^{-/-}$*; Il2rg*$^{-/-}$ mice following orthotopic transplantation with KPAR cells or *Etv4*$^{-/-}$ cells, n=5-9 per group.
(F) Heatmap showing hierarchical clustering of KPAR and *Etv4*$^{-/-}$ tumours based on mRNA expression of immune-related genes assessed by qPCR.
(G) Depletion of sgRNAs targeting Edn1 in WT versus *Rag2*$^{-/-}$*;Il2rg*$^{-/-}$ mice.
(H) Kaplan-Meier survival of immune-competent or *Rag2*$^{-/-}$*; Il2rg*$^{-/-}$ mice following orthotopic transplantation with KPAR cells or *Edn1*$^{-/-}$ cells, n=6-8 per group.
Data are mean ± SEM for (C-D and G). For (E and H), analysis of survival curves was carried out using log-rank (Mantel-Cox) test; * P<0.05, *** P<0.001.

#### Supp Figure 2

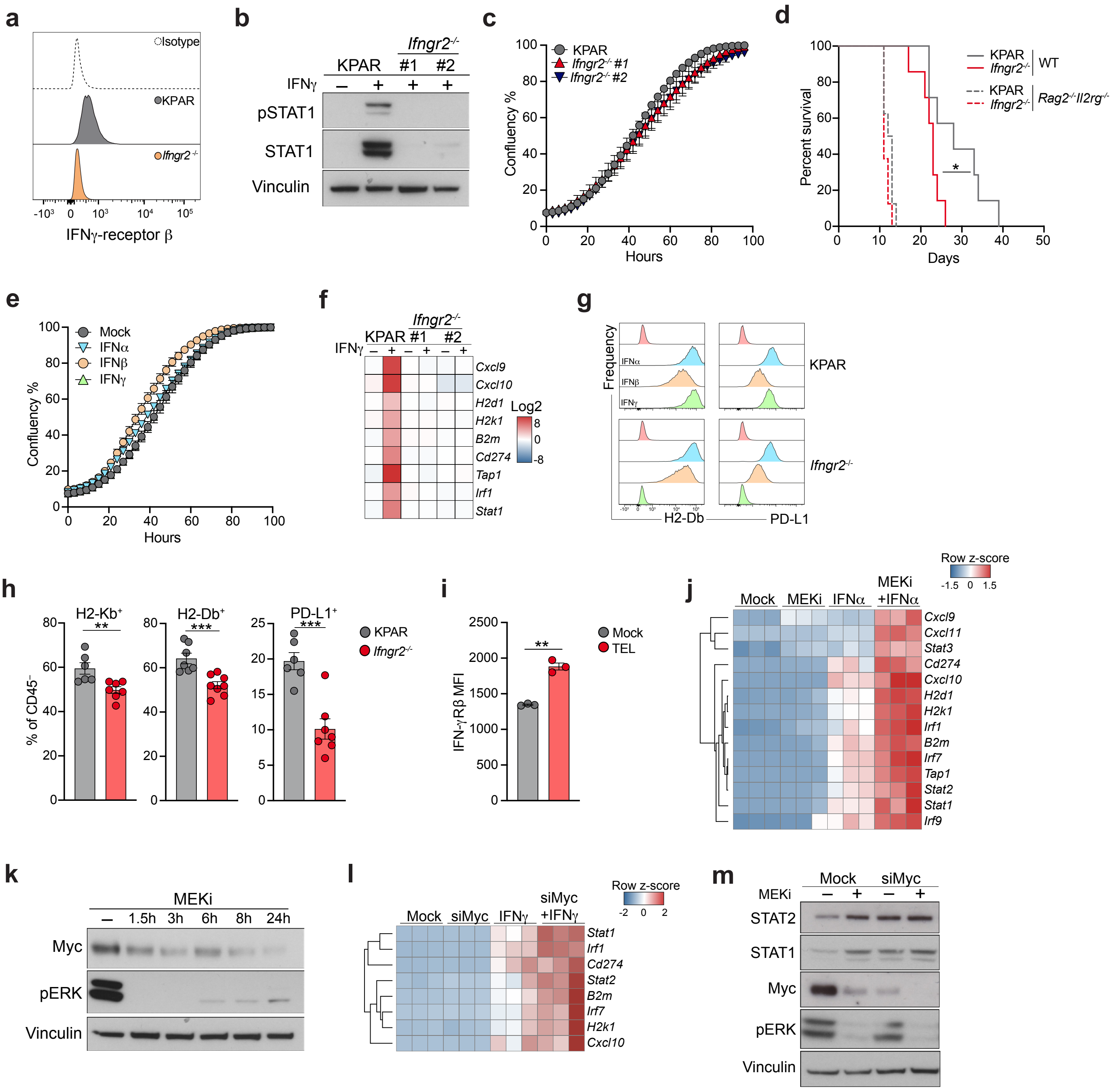


**Supplementary Figure 2. Oncogenic KRAS inhibits tumour-intrinsic IFN responses**

(A) Flow cytometry analysis showing surface expression of IFN-γ receptor subunit β on KPAR cells and *Ifngr2*-/- cells.
(B) Immunoblot for pSTAT1 and STAT1 in KPAR cells and *Ifngr2*-/- cells treated for 24h with 100ng/ml IFNγ.
(C) Incucyte analysis showing growth rate of KPAR cells and *Ifngr2*-/- cells in vitro.
(D) Kaplan-Meier survival of immune-competent or *Rag2*-/-; *Il2rg*-/- mice following orthotopic transplantation with KPAR cells or *Ifngr2*-/- (clone 2), n=5-10 per group. Analysis of survival curves was carried out using log-rank (Mantel-Cox) test; * P<0.05.
(E) Incucyte analysis showing growth of KPAR cells in the presence of 200ng/ml IFNα, 200ng/ml IFNβ or 100ng/ml IFNγ.
(F) Heatmap showing mRNA expression of IFN-response genes in KPAR cells and *Ifngr2*-/- cells treated with 100ng/ml IFNγ.
(G) Flow cytometry analysis showing surface expression of H2-Db (left) and PD-L1 (right) on KPAR cells and Ifngr2-/- cells treated for 24h with either 200ng/ml IFNα, 200ng/ml IFNβ or 100ng/ml IFNγ.
(H) Flow cytometry analysis showing frequency of H2-Kb, H2-Db and PD-L1 CD45- cells in KPAR and *Ifngr2*-/- tumours.
(I) Surface expression of the IFNγ-receptor β chain on KPAR cells treated with 10nM trametinib, 1μM linsitinib and 40nM everolimus for 24h.
(J) Heatmap showing expression of IFN-response genes by qPCR in KPAR cells treated for 24h with 200ng/ml recombinant IFNγ, 10nM trametinib, or both.
(K) Immunoblot for Myc in KPAR cells treated at indicated time points with 10nM trametinib.
(L) Heatmap showing expression of IFN-response genes in KPAR cells after siRNA-mediated knockdown of Myc and treatment with 100ng/ml IFN□.
(M) Immunoblot for STAT1 and STAT2 in KPAR cells after siRNA-mediated knockdown of Myc and treatment with 10nM trametinib for 24h. Data are mean ± SEM, for (C, E and H). Groups were compared using unpaired, two-tailed Student's t-test; ** P<0.01.

#### Supp Figure 3

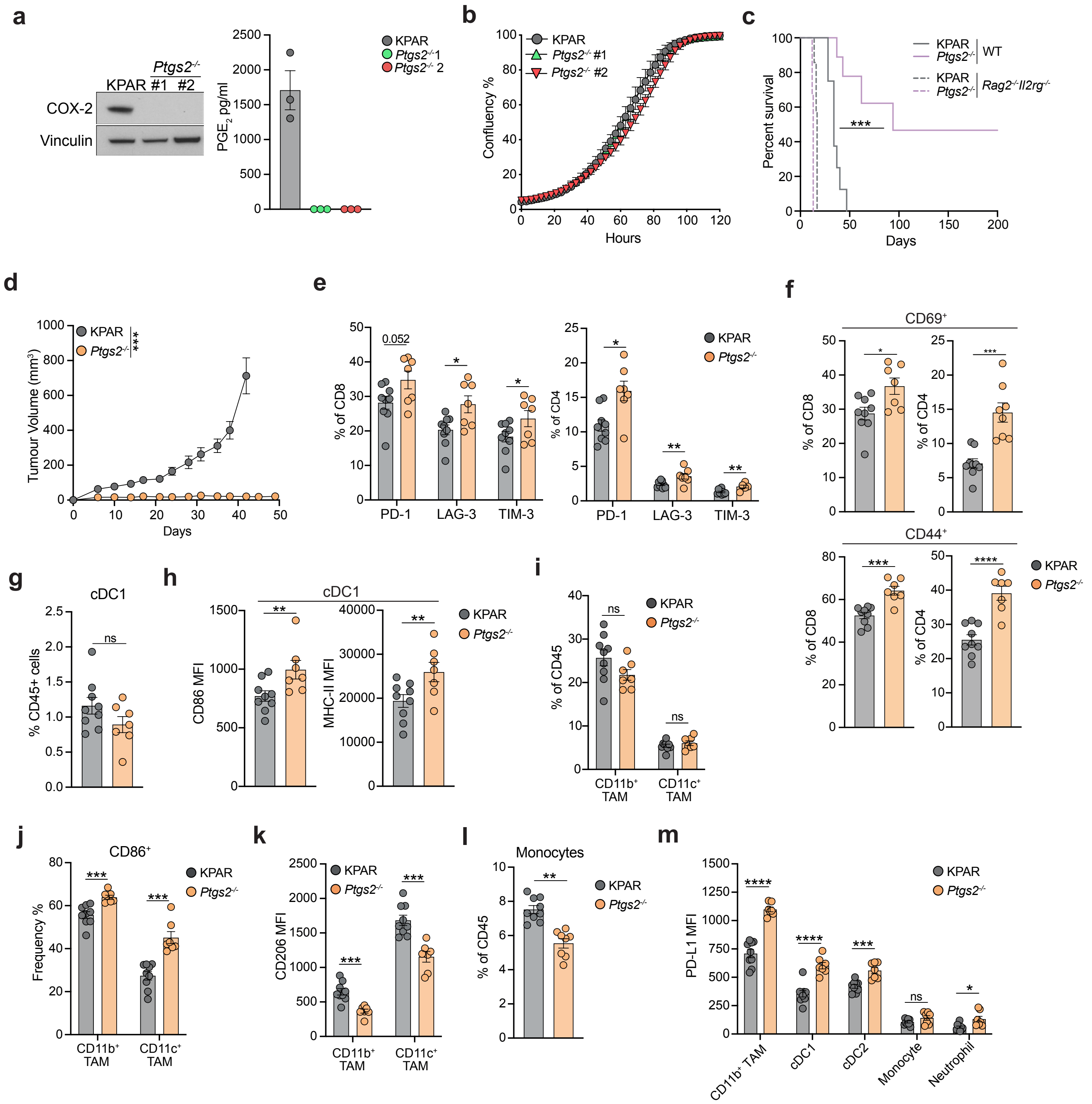


**Supplementary Figure 3. Tumour-intrinsic COX-2 remodels the lung tumour microenvironment**

(A) Immunoblot for COX-2 (left) and ELISA analysis for $PGE_2$ concentration (right) in KPAR cells and *Ptgs2*$^{-/-}$ cells.

(B) Incucyte analysis showing growth rate of KPAR cells and *Ptgs2*$^{-/-}$ cells *in vitro*.

(C) Kaplan-Meier survival of immune-competent or *Rag2*$^{-/-}$*; Il2rg*$^{-/-}$ mice following orthotopic transplantation with KPAR cells or *Ptgs2*$^{-/-}$ cells (clone 2), n=6-9 per group. Analysis of survival curves was carried out using log-rank (Mantel-Cox) test; *** P<0.001.

(D) Growth of subcutaneous KPAR and *Ptgs2*$^{-/-}$ tumours in immune-competent mice, n=10 per group.

(E) Quantification of PD-1, LAG-3 and TIM-3 surface expression on $CD8^+$ (left) and $CD4^+$ (right) T cells in KPAR and *Ptgs2*$^{-/-}$ tumours.

(F) Percentage of $CD69^+$ (left) and $CD44^+$ (right) T cells.

(G-H) Frequency of tumour-infiltrating cDC1s (F) and surface expression of CD86 and MHC-II on cDC1s (G).

(I-J) Frequency of tumour-infiltrating $CD11b^+$ and $CD11c^+$ TAMs (H) and percentage of $CD86^+$ TAMs (J).

(K) Surface expression of CD206 on TAMs.

(L) Frequency of $Ly6C^+$ monocytes.

(M) Surface expression of PD-L1 on myeloid cell populations.

For (A-B and E-M), data are mean ± SEM. For (E-M), n=7=9 per group. Samples were analysed using unpaired, two-tailed Student's t-test (E-M) or two-way ANOVA (D); ns, not significant, * P<0.05, ** P<0.01, *** P<0.001, **** P<0.0001.

#### Supp Figure 4

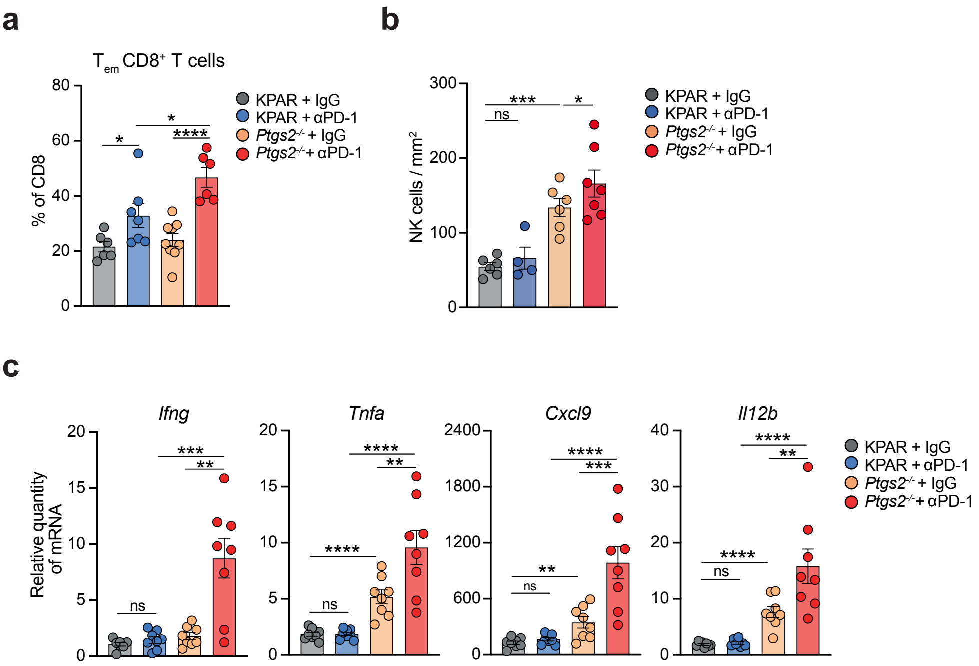


**Supplementary Figure 4. Genetic loss of COX-2 signalling synergises with ICB**
(A) Percentage of effector memory CD8+ T cells in KPAR or *Ptgs2*$^{-/-}$ tumours on day 7 after treatment with anti-PD-1 or corresponding isotype control (IgG).
(B) Quantification of CD8$^{+}$ T cells by immunohistochemistry in KPAR or *Ptgs2*$^{-/-}$ orthotopic tumours on day 7 after treatment with anti-PD-1 or corresponding isotype control (IgG).
(C) mRNA expression by qPCR of anti-tumour immunity genes in KPAR or *Ptgs2*$^{-/-}$ tumours treated as in (A).
Statistics were calculated using one-way ANOVA, FDR 0.05; ns, not significant, * P<0.05, ** P<0.01, *** P<0.001, **** P<0.0001.

#### Supp Figure 5

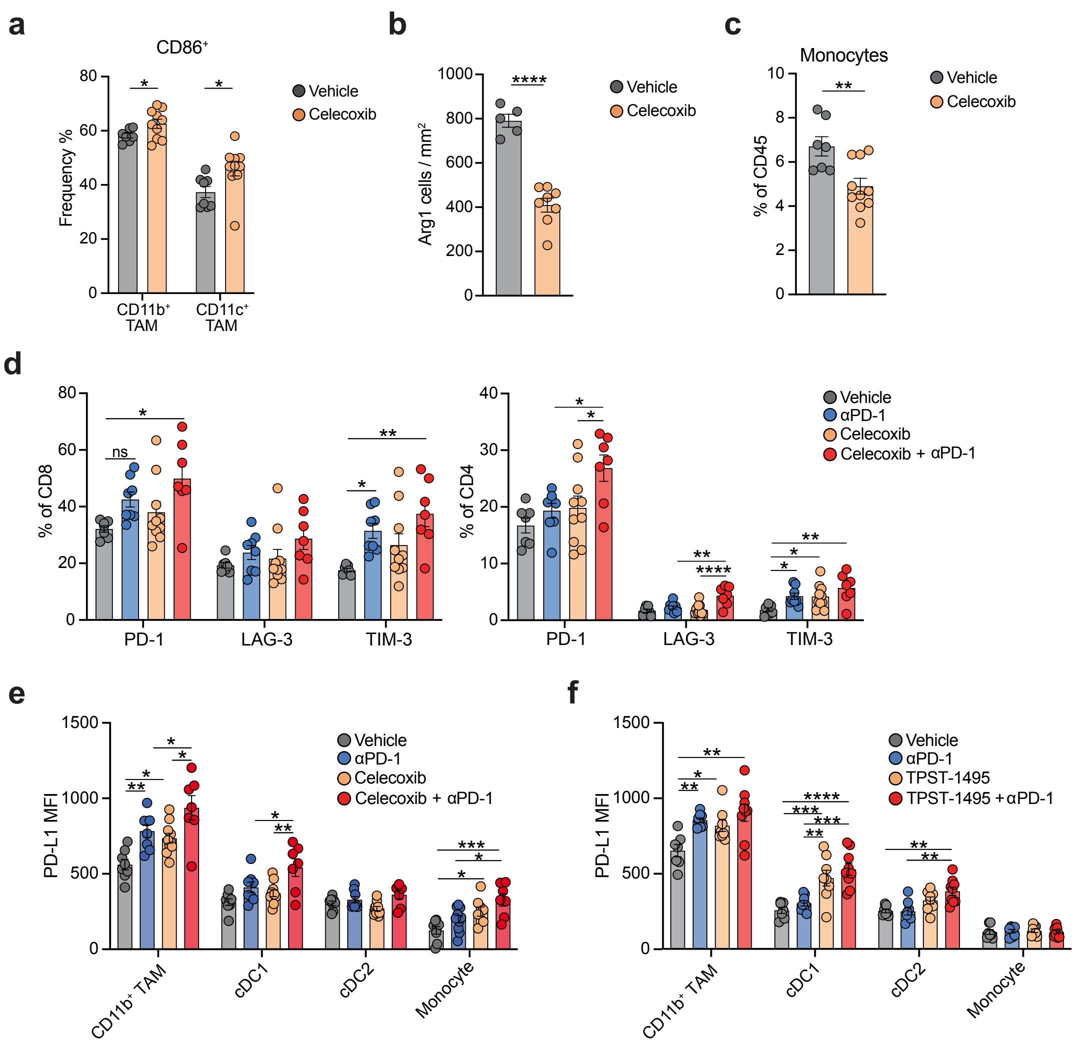


**Supplementary Figure 5. COX-2/PGE2 pathway inhibition remodels the TME and enhances the efficacy of ICB**
(A) Percentage of CD86⁺ TAMs in KPAR orthotopic tumours treated for 7d with 30mg/kg celecoxib.
(B) Quantification of Arg1⁺ cells by immunohistochemistry in KPAR orthotopic tumours treated as in (A).
(C) Frequency of Ly6C⁺ monocytes in KPAR tumours treated as in (A).
(D) Quantification of PD-1, LAG-3 and TIM-3 surface expression on CD8⁺ T cells (left) and CD4⁺ T cells (right) in KPAR tumours treated for 7d with celecoxib and/or anti-PD-1.
(E-F) Surface expression of PD-L1 on myeloid cell populations in KPAR tumours treated for 7d with anti-PD-1 and/or 30mg/kg celecoxib (E) or 100mg/kg TPST-1495 (F).
Data are mean ± SEM., n=5-10 per group. Groups were compared using unpaired, two-tailed Student's t-test (A-C) or one-way ANOVA, FDR 0.05 (D-F); ns, not significant, * P<0.05, ** P<0.01, *** P<0.001, **** P<0.0001.

### Supp Figure 6

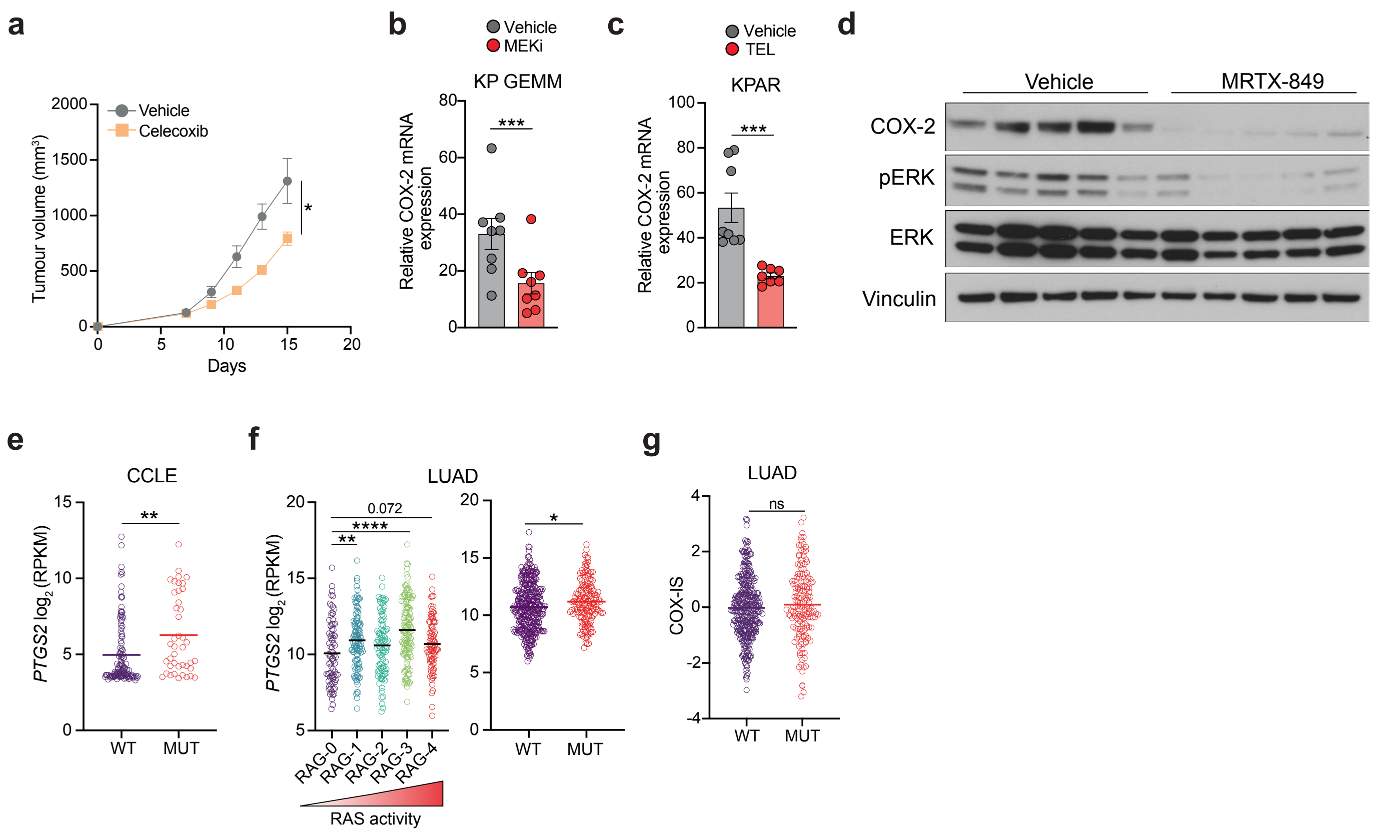


**Supplementary Figure 6. Oncogenic KRAS drives tumour-intrinsic expression of COX-2 in LUAD**

(A) Mean ± SEM 3LL ΔNRAS tumour volumes in mice treated with daily oral gavage of 30mg/kg celecoxib. Daily celecoxib treatment was initiated on day 7.

(B-C) COX-2 mRNA expression by qPCR in KP GEMM treated for 7d with 1.3mg/kg trametinib (B) or KPAR orthotopic tumours treated for 7d with 1.3mg/kg trametinib, 1.6mg/kg everolimus and 16.6mg/kg linsitinib (C).

(D) Immunoblot for COX-2 in KPAR$^{G12C}$ tumours treated for 7d with 50mg/kg MRTX849.

(E) COX-2 expression in RAS-WT and RAS-mutant human lung cancer cell lines from the CCLE database.

(F) COX-2 mRNA expression in LUAD samples from TCGA stratified by RAS-activity (left) or RAS mutational status (right).

(G) COX-IS in RAS-WT and RAS-mutant LUAD samples from TCGA.

For (A-C) data are mean ± SEM, n=7-9 per group. Samples were analysed using unpaired, two-tailed Student's t-test or one-way ANOVA, FDR 0.05; ns, not significant, * P<0.05, ** P<0.01, *** P<0.001, **** P<0.0001.

#### Supp Figure 7

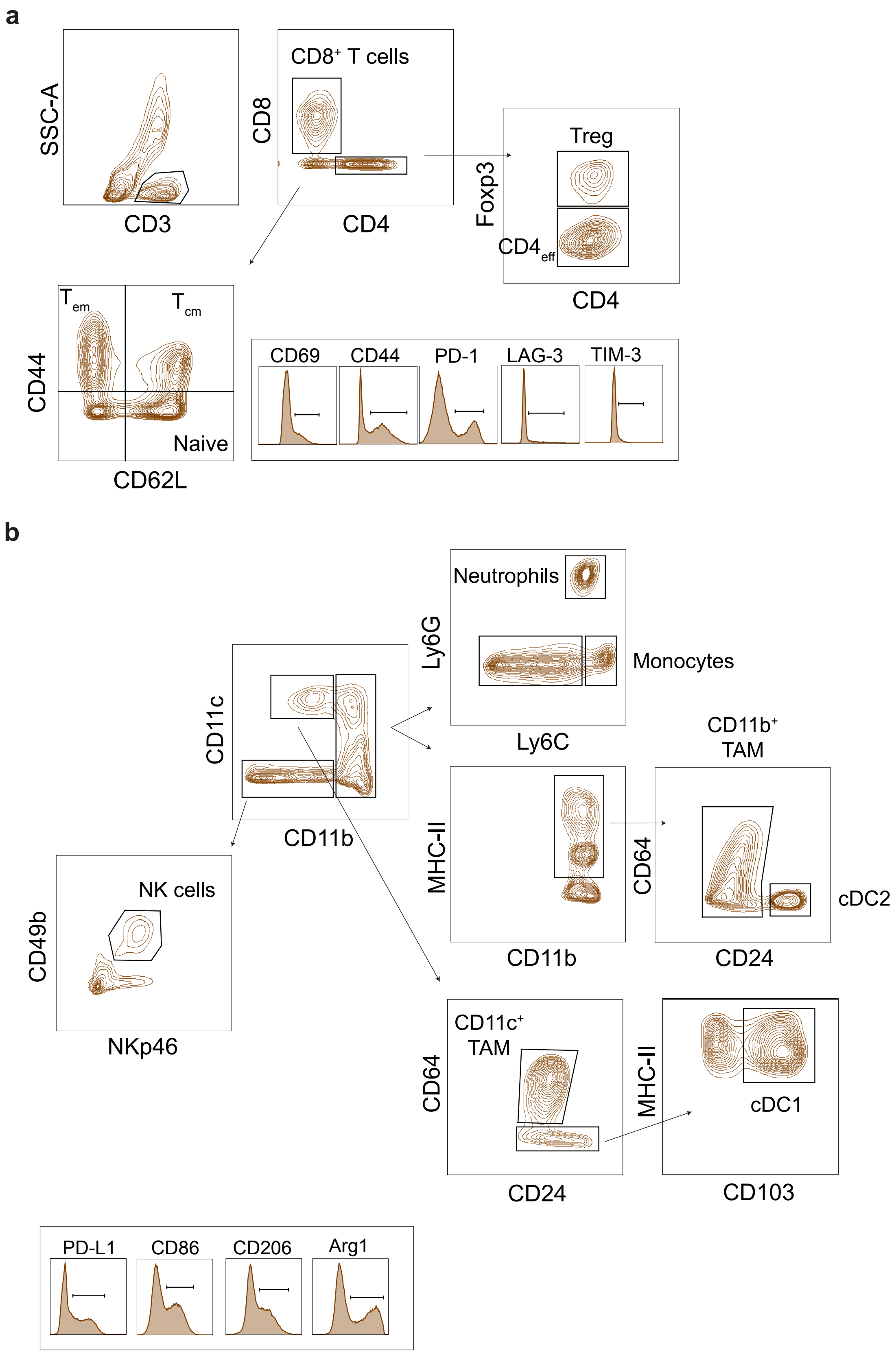


**Supplementary Figure 7. Flow cytometry gating strategies**

(A-B) Representative gating for the identification of different T cell subsets (A) and myeloid cells (B) and expression of phenotypic markers.
